# Spatially distinct ECM-producing fibroblasts and myonuclei orchestrate early adaptation to mechanical loading in the human muscle-tendon unit

**DOI:** 10.1101/2025.08.28.672815

**Authors:** Ask Møbjerg, Danielle Steffen, Peter Schjerling, Jens Rithamer Jakobsen, Anja Jokipii-Utzon, Mykhailo Y. Batiuk, Konstantin Khodosevich, Michael Rindom Krogsgaard, Valerio Izzi, Abigail L. Mackey, Michael Kjaer, Ching-Yan Chloé Yeung

## Abstract

Mechanical loading drives structural and functional improvements in muscle and tendon, protecting against injury at their interface – the myotendinous junction (MTJ) – and within the tendon matrix. However, the early cellular and molecular events that initiate these adaptations in humans remain poorly understood. To investigate this, we applied single nucleus RNA sequencing and in situ hybridization to map the acute transcriptional response of the human muscle-tendon unit to a single bout of eccentric resistance exercise, with a focus on extracellular matrix (ECM) regulation. We identified four transcriptionally distinct fibroblast subtypes expressing key ECM components, including *COL1A1* and *DCN*. Three of these subtypes were localized to tendon and responded to exercise: two were spatially restricted to the collagen fascicles or the MTJ, while the third, enriched in the interfascicular matrix (IFM), exhibited the strongest response. This IFM population, marked by *PDGFRA*, upregulated *PRG4* and *VCAN*, ECM genes linked to tissue lubrication and resilience. In parallel, exercise induced dynamic ECM regulation in myonuclei, particularly in a distinct subset of type II myonuclei at the MTJ that expanded in number and robustly upregulated *COL22A1*, a collagen essential for MTJ integrity. Together, these findings uncover a spatially organized, cell type-specific program of ECM remodeling in response to mechanical load, offering new insight into the early molecular events of human muscle-tendon adaptation.

## INTRODUCTION

The muscle-tendon unit is essential for force transmission in the movement of the human body, and its function relies on the homeostasis of the extracellular matrix (ECM). Further, tendons account for over 30% of musculoskeletal injuries^1^, with the myotendinous junction (MTJ) being the most injury-prone region to acute damage^2^. Mechanical loading, especially resistance training involving eccentric contractions, strengthens both muscle and tendon and reduces the overall risk for strain injuries^3–6^. Yet, the early molecular events that initiate these protective adaptations remain poorly understood. Because long-term tissue remodeling is initiated by acute transcriptional responses, identifying the immediate cellular and molecular changes following mechanical load is essential for understanding how tissue resilience (structural integrity and injury resistance) is built and how injuries might be prevented. However, it is not known which cell types initiate these changes, nor how ECM remodeling is regionally and temporally regulated across the muscle-tendon unit.

The molecular effects of resistance training have been more thoroughly investigated in skeletal muscle, where transcriptional and metabolic responses to various training stimuli have been mapped^7,8^. In contrast, much less is known about the MTJ, a specialized interface enriched in collagen XXII and defined by *COL22A1* expression in both MTJ-localized myonuclei and adjacent mononuclear cells (tenocytes)^9,10^. A previous study revealed the MTJ region upregulates key ECM molecules known to be enriched in the MTJ^11^ to a single bout of eccentric training, measured 7 days post-exercise^12^. However, immunofluorescence-based analyses found no changes in major collagens (types I, III, VI, XII, XIV, XXII) at the MTJ either 2-3 days after a single training session or following 4 weeks of repeated eccentric exercise, in contrast to detectable changes within the adjacent intramuscular connective tissue^13^. Together, these findings suggest that MTJ adaptation may occur through mechanisms not captured by protein abundance alone, and that the identity and behavior of the specific cell populations orchestrating this response remain poorly defined.

Tendon, on the other hand, is relatively quiescent and remodeling of its collagen-rich ECM is typically observed in response to chronic overuse or injury^14,15^. However, a short-term stable isotope study showed that tendon has a basal protein synthesis rate comparable to muscle^16^, suggesting a capacity for acute adaptation beyond collagen production. Indeed, no change in peritendinous collagen synthesis was detected after 12 weeks of eccentric training in healthy Achilles tendons using microdialysis, with increases only observed in chronically overloaded tissue^17^. Nonetheless, other studies have shown that high-load eccentric training improves the mechanical stiffness and load-bearing capacity in healthy patellar tendons^18,19^, implying that tendon adaptation may involve structural or cellular changes not captured by collagen synthesis alone.

Recent studies also challenge the notion of tendon fibroblasts as a homogeneous population. Human tendons are composed of aligned collagen fascicles housing elongated tenocytes and an interfascicular matrix (IFM) containing rounded fibroblasts within an ECM niche enriched in elastin and proteoglycans such as lubricin and versican^20^. The IFM enables fascicle sliding during loading, a function that declines with age^21^. Single cell RNA sequencing of aging equine tendon recently revealed multiple IFM fibroblast subtypes^22^, and single nucleus RNA sequencing (snRNAseq) of human hamstring tendon identified additional collagen-producing cell types beyond classical fibroblasts, including satellite cells, mural cells, and adipocytes^23^. These findings highlight the cellular diversity of the muscle-tendon unit and underscore the need to resolve which resident cell types are transcriptionally responsive to mechanical loading.

Here, we applied snRNAseq to hamstring muscle-tendon tissues collected 4 h after a single session of eccentric resistance exercise in healthy adults. Our aim was to map early, cell type-specific transcriptional responses, focusing on cell types that regulate the ECM.

## RESULTS

### Single nucleus transcriptomic profiling of human hamstring tissues

Twelve healthy individuals participated but due to insufficient nuclei (n=1 control and n=1 exercised) and sequencing quality issues (n=1 control and n=1 exercised) (see Materials and Methods), samples from 8 individuals were analyzed (n=4 control, n=4 exercised; Table 1). No significant differences were found in the age, but there was an uneven sex distribution, with four males in the control group and three females and one male in the exercise group (Table 1). We profiled the transcriptomes of 61,251 nuclei from hamstring tendon (either semitendinosus or gracilis; see Table 1): 26,524 control and 34,691 exercised. The most abundant transcript identified was *MALAT1*, a nuclear transcript^24^, confirming that our data set contained nuclear RNAs.

**Table 1.** Age and sex distribution of samples. Mean ± SD are shown. No significant differences were found between the age of control and exercised subjects (p=0.89 for included and p=0.71 for final samples analyzed; t-tests). No significant differences were found between the tissue weight of control and exercised subjects (p=0.30 for included and p=0.26 for final samples analyzed; t-tests).

|  | Control | Exercised |
| --- | --- | --- |
| *Included* |  |  |
| **N** | 6 | 6 |
| **Age, years (range)** | 27.0 ± 8.6 (21-42) | 27.8 ± 9.9 (19-45) |
| **Sex** | 2 F, 4M | 3 F, 3 M |
| **Tissue type** | 4 gracilis, 1 semitendinosus | 2 gracilis, 2 semitendinosus,<br>1 mixed |
| **Tissue weight (mg)** | 380 ± 110 (186-491) | 309 ± 113 (119-407) |
| *Final analysis* |  |  |
| **N** | 4 | 4 |
| **Age, years (range)** | 29.3 ± 10.2 (21-42) | 32.0 ± 9.6 (22-45) |
| **Sex** | 4 M | 3 F, 1 M |
| **Tissue type** | 3 gracilis, 1 semitendinosus | 1 gracilis, 2 semitendinosus,<br>1 mixed |
| **Tissue weight (mg)** | 376 ± 132 (186-491) | 267 ± 118 (119-407) |

Initial clustering analysis using Seurat identified 18 nuclei populations, which, based on conserved marker genes, were assigned to seven major cell groups: immune (2 clusters), adipocyte, endothelial (2 clusters), smooth muscle (2 clusters), fibroblasts (4 clusters), satellite cells and myonuclei (6 clusters including MTJ) (Fig. 1A and Data S1). All cell populations were represented across all samples (Fig. S1A-B). Uniform Manifold Approximation and Projection (UMAP) embedding was used for visualization, showing a clear separation between myonuclei and non-myonuclei, with *TTN* expression confirming the identity of the muscle-derived cluster (Fig. S1C). Stratifying the data by biological sex and exercise did not reveal major differences in the representation of nuclei in individual clusters (Fig. S1D-E). Expression of selected marker genes revealed largely distinct transcriptomic profiles across clusters (Fig. 1B). Myonuclei clusters shared highly similar signatures, all marked by strong expression of *TTN*, *ACTA1*, *NCAM1*, and fiber type-specific myosins (*MYH1*, *MYH2*, *MYH7*), genes known to be expressed by myonuclei in the MTJ region of the hamstring^25^ (Fig. 1B). Notably, *COL22A1* expression was almost exclusively restricted to MTJ myonuclei, with some additional expression observed in one fibroblast cluster (Fib1), suggesting these populations are associated closest to the MTJ^25^ (Fig. 1B). Satellite cells were identified by the expression of *PAX7*, the transcription factor that regulates myogenesis^26^. The expression of early B-cell factor-1 (*EBF1*) was present in all non-myonuclei, except immune cell clusters. All four fibroblast clusters expressed ECM genes, *COL1A1*, *COL3A1, COL6A1* and *DCN* (Fig. 1B). Smooth muscle cell clusters were defined by *ACTA2* and *FHL5* expression. Endothelial cells expressed *PECAM1* and *VWF*. Adipocytes expressed the adipogenic markers *PLIN1* (perilipin 1) and *PPARG*. Immune and T cell clusters expressed *DOCK2*, with the immune cluster uniquely expressing *F13A1* and the macrophage marker *CD163*, while T cells were marked by *CD247* and *SKAP1* expression (Fig. 1B).

**Figure 1.**
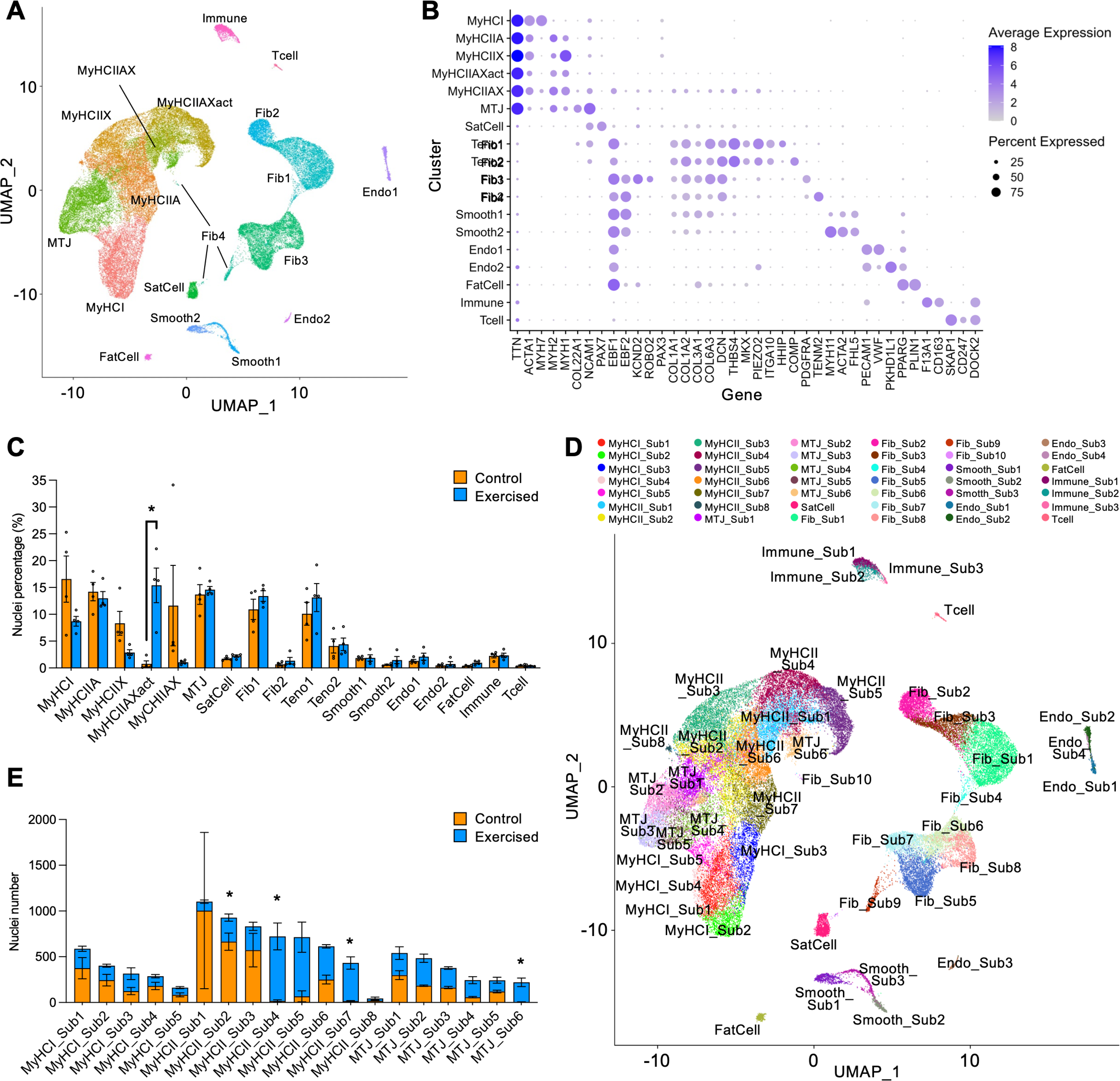
Single nucleus RNA-seq analysis of human hamstring tendons following a single bout of eccentric resistance exercise. **(A)** Uniform manifold approximation and projection (UMAP) embedding of 61,215 nuclei from healthy hamstring tendons of 8 individuals (control and exercised combined) showing 18 clusters belonging to seven cell types: myonuclei (including MTJ myonuclei), satellite cells, fibroblasts, smooth muscle cells, endothelial cells, fat cells, and immune cells. **(B)** Dot plot of selected markers that define each nuclei cluster. Expression values displayed are derived from RNA assay data obtained from individual nuclei. **(C)** Changes in nuclei proportion in clusters with exercise (*FDR<0.05, DESeq (Wald test with Benjamini-Hochberg correction)). Bars show mean ± SEM. **(D)** UMAP showing nuclei subclusters in healthy human hamstring tendons. **(E)** Changes in myonuclei proportion in myonuclei subclusters (*FDR<0.05, DESeq (Wald test with Benjamini-Hochberg correction)). Bars show mean ± SEM.

### MTJ myonuclei changes in response to exercise

With exercise, a significant increase of ∼16-fold in nuclei abundance was observed in the MyHCIIAX-activated (MyHCIIAXact) cluster, characterized by co-expression of *MYH1* and *MYH2*, encoding type IIX and IIA myosin heavy chain, respectively (FDR=6.6 × 10^−3^, DESeq (Wald test with Benjamini-Hochberg correction); Fig. 1C, S1E and S2). This change is consistent with the activation of type II muscle fibers by the intended resistance training stimulus^27,28^. To delve into the differences with exercise, we performed subclustering of the 18 main clusters, resulting in 42 transcriptionally distinct subpopulations (Fig. 1D and Data S2). At the subcluster level, significant changes in nuclei abundance with exercise were observed in 4 subclusters, all type II myonuclei (FDR<0.05, DESeq (Wald test with Benjamini-Hochberg correction); Fig. 1E). MTJ_Sub6, one of the MTJ subclusters, exhibited a substantial response to exercise, with ∼97% of nuclei derived from the exercised group (FDR=3.2 × 10^−14^, DESeq (Wald test with Benjamini-Hochberg correction); Fig. 1E). Similarly, ∼98% of MyHCII_Sub4 and ∼96% of MyHCII_Sub7 myonuclei were derived from the exercised samples (FDR=4.0 × 10^−6^ and 5.6 × 10^−12^, respectively; DESeq (Wald test with Benjamini-Hochberg correction); Fig. 1E). Conversely, ∼72% of MyHCII_Sub2 nuclei were derived from the control group (FDR=1.0 × 10^−5^; DESeq (Wald test with Benjamini-Hochberg correction); Fig. 1E). Gene ontology (GO) enrichment analysis of differentially expressed genes (DEGs) in the four exercise-responsive myonuclei subclusters revealed subcluster-specific programs related to cytoskeletal organization, protein folding, metabolic regulation, and myofibrillar remodeling (Data S3 and Data S4). These findings indicate that type II myonuclei engage distinct stress-adaptive and structural transcriptional responses following exercise.

### Fibroblast diversity in human hamstring muscle-tendon unit

Four fibroblast clusters, comprising 17,863 nuclei, were identified. Fib3 comprised the largest proportion (42.4 ± 2.7%, mean ± SEM), closely followed by Fib1 (39.6 ± 3.8%), Fib2 (14.4 ± 2.3%) and Fib4 (3.5 ± 1.0%) (Fig. 2A-B). We performed differential expression analysis across the 18 main clusters to identify marker genes enriched in each of the four fibroblast populations. This analysis revealed 52, 47, 56, and 48 uniquely enriched genes in Fib1, Fib2, Fib3, and Fib4, respectively, with minimal overlap between clusters (Fig. 2C and Data S1). Matrisome analysis revealed that a total of 38 core matrisome and matrisome-associated genes among these fibroblast marker genes (Fig. S3). Fib1 and Fib2 expressed high levels of *COL1A2*, *THBS4*, and also expressed the transcription factor mohawk (*MKX*), which controls tenogenic differentiation^29,30^ (Fig. 2D). Fib1 contained high levels of *PIEZO2*, a mechanosensitive ion channel activated during tendon loading^31^, and uniquely expressed *HHIP* and high levels of *ITGA10*, which are highly enriched at the MTJ^11,32^ (Fig. 2E). When comparing marker gene expression among the four fibroblast clusters, the classic MTJ marker *COL22A1* was significantly upregulated in Fib1 (Fig. 2E and Data S5). Fib2 nuclei exclusively expressed *COMP*, a glycoprotein produced in tendon and is upregulated with load^33^, supporting their identity as the main population of fibroblasts in the tendon fascicular matrix (Fig. 2F). Fib3 nuclei were defined by the unique expression of *PDGFRA* and *SCARA5*, as well as high expression of ECM markers identifying IFM cells^22^, *COL14A1*, *TNXB* and *VCAN* (Fig. 2G and S3). Fib4 nuclei had the unique expression of *TENM2*, *PAK3*, *ITGA6* and *SLC22A3* (Fig. 2H).

**Figure 2.**
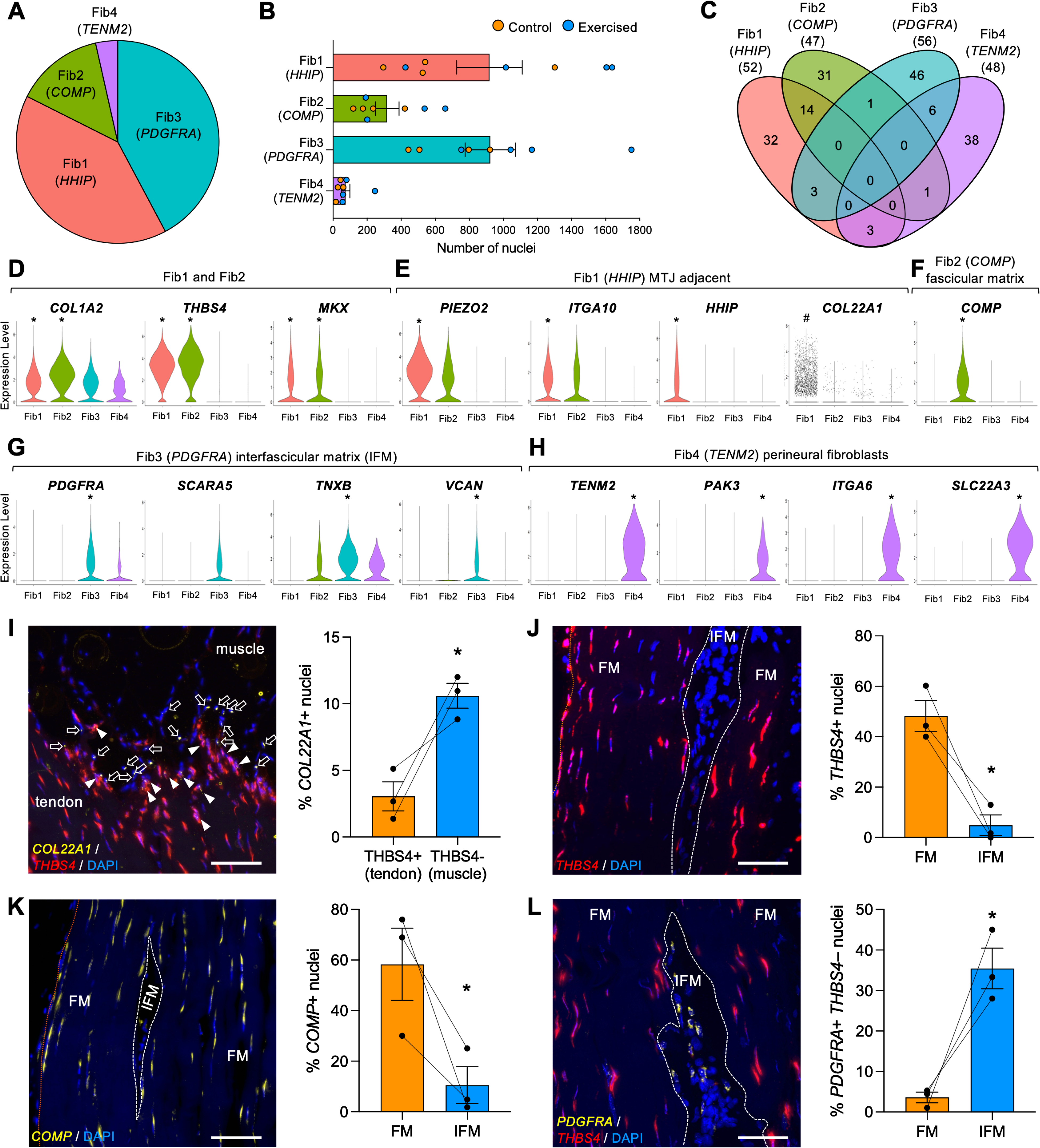
Distinct fibroblast populations in healthy human hamstring tendons and their spatial localization. **(A)** Proportion of fibroblast nuclei in healthy human hamstring tendons. Fib1, 39.6 ± 3.8% (mean ± SEM); Fib2, 14.4 ± 2.3%; Fib3, 42.4 ± 2.7%; Fib4, 3.5 ± 1.0% (n=8). **(B)** Number of nuclei from each sample for the 4 fibroblast clusters (n=8). Bars show mean ± SEM. **(C)** Venn diagram showing minimal overlap in DEGs among the four fibroblast clusters. **(D-H)** Violin plots showing the expression of selected marker genes that distinguish between the tendon fibroblasts clusters, **(D)** Fib1 and Fib 2: *COL1A2*, *THBS4*, and *MKX*, **(E)** Fib1 MTJ adjacent fibroblasts: *PIEZO2*, *ITGA10*, *HHIP*, and when tested within fibroblasts subclusters, *COL22A1* (with individual points overlaid), **(F)** Fib2 fascicular matrix fibroblasts: *COMP*, **(G)** Fib3 interfascicular matrix fibroblasts: *PDGFRA*, *SCARA5*, *TNXB*, and *VCAN*, and **(H)** Fib4 perineural fibroblasts: *TENM2*, *PAK3*, *ITGA6*, and *SLC22A3*. * indicates significance (log2 fold change>1, FDR<0.05, DESeq) when compared to all clusters. ^#^ indicates significance (log2 fold change>1, FDR<0.05, DESeq) when compared within the fibroblast subclusters. **(I-L)** Quantification of RNAscope staining for nuclei with specific transcripts: **(I)** *COL22A1*, **(J)** *THBS4*, **(K)** *COMP*, and **(L)** *PGDFRA* in selected regions of longitudinally sectioned hamstring tissues (n=3). Scale bars, 100 µm. White arrowheads indicate *COL222A1+ THBS4+* nuclei. White dashed lines indicate the boundaries between the IFM and fascicular matrix. Orange dotted lines indicate the borders of the tendon tissue. FM, fascicular matrix. IFM, interfascicular matrix (encompassed by the dotted white line). Bars show mean ± SEM. Percentages are calculated from the following total number of nuclei per sample: **(I)** 150, 219, 215, **(J)** 214, 256, 90, **(K)** 152, 95, 107, and **(L)** 120, 150, 96. White open arrows indicate *COL222A1+ THBS4−* nuclei. * indicates significance (p=9.8 × 10^−4^ for *COL22A1*, p=4.7 × 10^−14^ for *THBS4*, p=2.0 × 10^−12^ for *COMP*, and p=1.3 × 10^−10^ for *PDGFRA*; from mixed-effects logistic regression). Donor age and sex information for RNAscope samples are provided in Data S12.

To spatially validate the identity and anatomical context of the transcriptionally defined fibroblast populations, we performed RNAscope in situ hybridization on human tendon sections. Due to the tendon-optimized pretreatment protocol, muscle cytoplasm was largely cleared by the protease treatment, but myonuclei remained intact (Fig. S4A-B). Staining patterns varied depending on the target RNA, reflecting differences in transcript abundance and subcellular localization^34^. Fib1 marker *COL22A1* was mostly detected in *THBS4−* nuclei located in the border between tendon and muscle, which indicated MTJ myonuclei^9^ (Fig. 2I). Consistent with the snRNAseq data, in which only ∼24% of Fib1 nuclei expressed *COL22A1* and at lower levels than MTJ myonuclei, we observed very few nuclei co-expressing *COL22A1* and *THBS4*, and these were confined to the MTJ region (Fig. 2I). *THBS4*, expressed by Fib1 and Fib2, was broadly distributed throughout the tendon body and showed significantly reduced expression in IFM regions (Fig. 2J). *COMP*, a unique marker to Fib2, was similarly restricted to the collagen fascicles (Fig. 2K). The Fib3 marker *PDGFRA* was significantly enriched in the IFM, validating the transcriptional annotation and resolving the cellular identity of this fibroblast subtype (Fig. 2L). Attempts to localize Fib4 to tendon using *TENM2* were unsuccessful (Fig. S4C), possibly owing to its low abundance.

To place these transcriptionally defined fibroblast populations in the context of existing tendon datasets, we performed cross-study integration with published snRNAseq data from human tendon and MTJ tissues. As expected, our dataset showed great similarity to a human hamstring study^23^. The integration revealed that Fib1 and Fib2 tendon fascicle fibroblasts aligned with *“MKX⁺ fibroblasts”*, while Fib3 IFM fibroblasts corresponded to *“PDGFRA⁺ fibroblasts”*, the location of which was previously predicted to be near skeletal muscle using spatial transcriptomics data, and Fib4 corresponded to *“nerve cell”*^23^ (Fig. S5A). When integrated with the human hamstring MTJ dataset^25^, all myonuclei clusters from the present study correlated with *“MTJ”*. In contrast, integration with the quadriceps tendon dataset^35^ revealed only partial overlap with our fibroblast clusters (Fig. S5B-C). Fib4 corresponded to *“nerve cells”* in the human hamstring dataset but also retained a non-tenocyte *“fibroblast”* identity in the MTJ dataset (Fig. S5A-B). To investigate whether Fib4 may correspond to a skeletal muscle-resident fibroblast, we integrated with a human skeletal muscle aging atlas^36^ and found high correlation with *“perineural fibroblasts”* (Fig. S5D). In line with this identity, Fib4 uniquely expressed *ITGA6* and *SLC22A3*, markers identifying perineural fibroblasts^36^, which could explain the mix of nerve cell and fibroblast identities, as well as its low nuclei abundance in our samples and absence in tendon sections (Fig. 2H). RNAscope experiments to localize Fib4 to MTJ-adjacent muscle using *TENM2* remained inconclusive due to the low frequency of *TENM2*-positive nuclei (Fig. S4E).

### Subclustering reveals fibroblast subpopulations with distinct exercise responses

We next applied DESeq2 to pseudobulked data from the four fibroblast clusters to assess their acute transcriptional response to exercise (Data S6). Only Fib1-3 showed statistically significant changes to gene expression (Fig. S6A). There was a modest shift in gene expression 4 hours after a single bout of heavy resistance exercise in the fibroblasts, with 50, 23 and 115 DEGs in each fibroblast cluster, respectively (Fig. S6A and Data S6). While Fib1 and Fib2 DEGs were mostly downregulated, Fib3 exhibited 45 upregulated DEGs, including the proteoglycans *PRG4* and *VCAN* (Fig. S6A). Six DEGs were shared across the three fibroblast clusters (Fig. S6B). Of these, five were downregulated, including the co-chaperones *CHORDC1* and *DNAJB6* (Fig. S6B and C). *XIST*, a long non-coding RNA produced by the X chromosome, showed higher expression in the exercise group due to the sex imbalance in the control and exercised groups (Fig. S6C). STRING analysis revealed potential interactions between the DEGs (Fig. S6D). Significant functional enrichment was observed only in the DEG networks of Fib1 and Fib3. In Fib1, DEGs were enriched for terms related to the regulation of biological quality and the autolysosome cellular compartment (Data S7). In Fib3, DEGs were enriched for pathways associated with angiogenesis, tissue development, and the extracellular matrix compartment (Data S7).

Given the modest cluster-level changes with exercise and known fibroblast heterogeneity, we examined the fibroblast subclusters (Data S2), which identified ten transcriptionally distinct subpopulations (Fig. 3A-B and Table 2). Fib_Sub1 and Fib_Sub3 differentially express the MTJ marker, *HHIP,* and high levels of *PIEZO2* (Fig. S7A-B and Data S2). When tested within the fibroblast subcluster groups, Fib_Sub1 expressed high levels of MTJ ECM proteins, *COL22A1* and *POSTN*, consistent with a localization near the muscle-tendon junction (Fig. S7B and Data S8). Fib_Sub2 exclusively expressed COMP, identifying them as fascicular matrix fibroblasts (Fig. S7C). These three subtypes of fascicular matrix fibroblasts responded to exercise similarly, with GO analysis revealing pathways related to chaperone activity, protein folding and stress response (Fig. 3C and Data S4). Fib_Sub4 comprised relatively few nuclei (34 ± 35, mean ± SD), showed a mixed transcriptional profile of fascicular and IFM fibroblasts, and exhibited downregulation of only two genes in response to exercise (Fig. 3A-C and S7D). Fib_Sub5-8 commonly expressed high levels of *KCND2*, *PDGFRA*, *NEGR1* and *SCARA5*. And consistent with an IFM fibroblast identity, there was high expression of the aforementioned markers^22^, *TNXB* and *VCAN* (Fig. S7E and Data S8). GO analysis, however, revealed divergent responses to exercise; Fib_Sub5 responded similarly to Fib_Sub1-4, by regulating chaperone and stress response pathways (Fig. 3C and Data S9). Fib_Sub6 exhibited the most extensive response, with 167 DEGs belonging to vascular remodeling, tissue morphogenesis, and matrix adhesion pathways (Fig. 3C and Data S9). GO analysis of the small 20-DEG set in Fib_Sub7 suggested potential involvement in energy metabolism (Fig. 3C and Data S9). Fib_Sub8 responded to exercise by regulating genes related to broad developmental processes (Fig. 3C and Data S9). Fib_Sub9 expressed *TENM2*, defining them as the perineural fibroblasts, while Fib_Sub10 was spatially distinct from other fibroblast clusters on the UMAP and characterized by high expression of *NRXN1* and *GRIK2* (Fig. 3A and S7F-G). Neither cluster showed significant transcriptional changes in response to exercise (Table 2).

**Figure 3.**
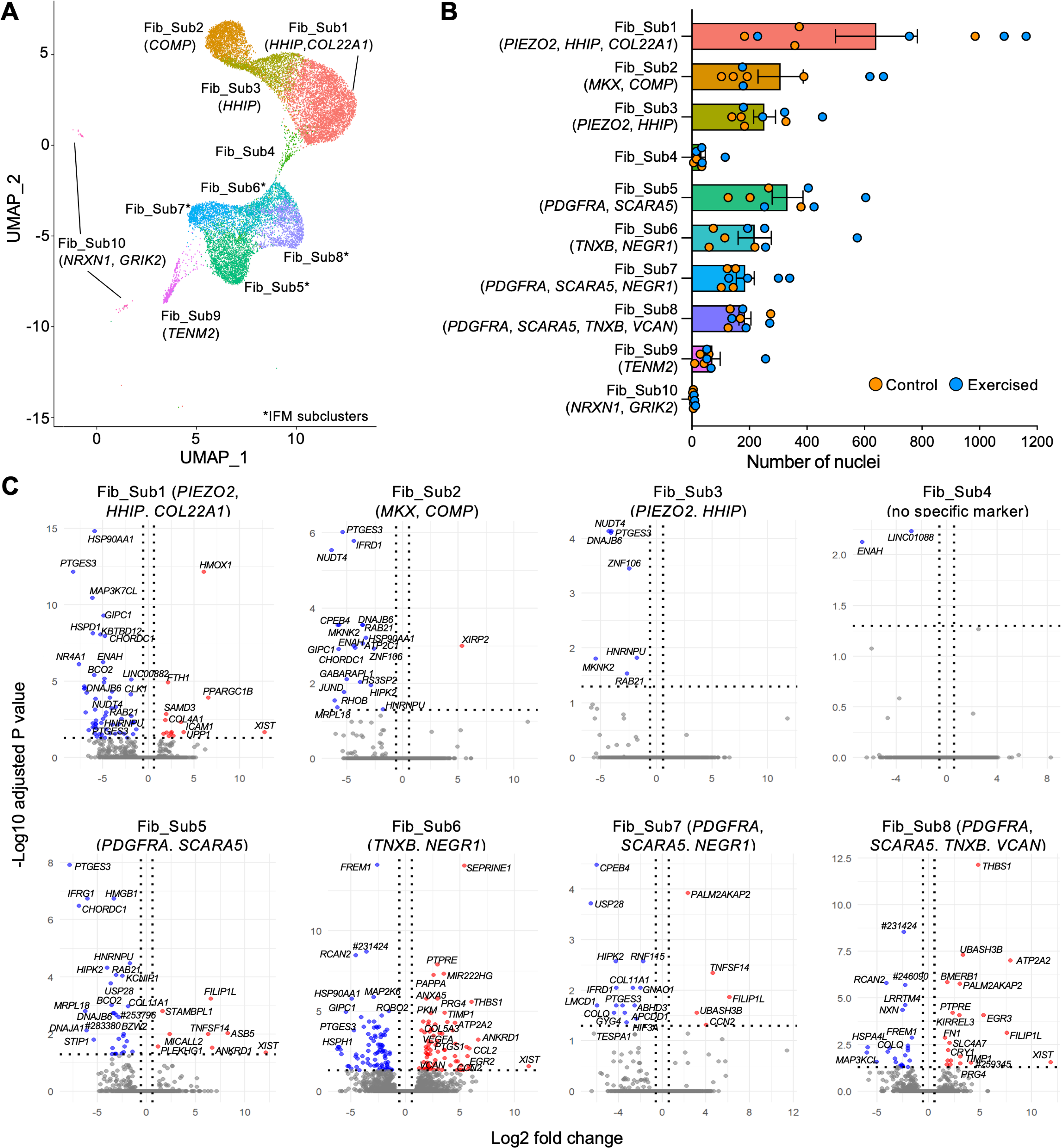
Exercise regulated RNAs in human hamstring tendon fibroblasts. **(A)** UMAP showing the fibroblast subclusters, Fib_Sub1-10. * indicates the IFM subclusters: Fib_Sub5, Fib_Sub6, Fib_Sub7 and Fib_Sub8. **(B)** Number of nuclei from each sample for the 10 fibroblast subclusters (n=8). Bars show mean ± SEM. **(C)** Volcano plots showing exercise-induced DEGs in Fib_Sub1-8. No exercise-induced DEGs were detected in Fib_Sub9 or Fib_Sub10. Blue data points indicate significantly downregulated DEGs by >-0.585 log2 fold change (FDR<0.05, DESeq2). Red data points indicate significantly upregulated DEGs by >0.585 log2 fold change (FDR<0.05, DESeq2).

**Table 2.** Number of exercise-induced DEGs in human hamstring subclusters. Total number of DEGs, including protein-coding and long non-coding RNAs. A dashed line indicates no DEGs detected at all. Numbers in brackets indicate number of matrisome-annotated DEGs (including secreted factors). For full lists of exercise-induced DEGs and matrisome-annotated DEGs, see Data S3 and S10, respectively.

| Sub cluster ID | Total exercise DEGs | Upregulated | Downregulated |
| --- | --- | --- | --- |
| MyHCI_Sub1 | 427 (25) | 286 (20) | 141 (5) |
| MyHCI_Sub2 | 152 (8) | 103 (4) | 49 (4) |
| MyHCI_Sub3 | 1365 (55) | 755 (38) | 610 (17) |
| MyHCI_Sub4 | 439 (24) | 276 (18) | 163 (6) |
| MyHCI_Sub5 | 362 (25) | 256 (22) | 106 (3) |
| MyHCII_Sub1 | 887 (43) | 711 (41) | 176 (2) |
| MyHCII_Sub2 | 2717 (96) | 1607 (82) | 1110 (14) |
| MyHCII_Sub3 | 2305 (98) | 1771 (92) | 534 (7) |
| MyHCII_Sub4 | 1155 (39) | 820 (35) | 335 (4) |
| MyHCII_Sub5 | 981 (29) | 815 (26) | 166 (3) |
| MyHCII_Sub6 | 4778 (145) | 2450 (123) | 2328 (22) |
| MyHCII_Sub7 | 1430 (44) | 835 (37) | 585 (7) |
| MyHCII_Sub8 | - | - | - |
| MTJ_Sub1 | 3103 (103) | 1788 (92) | 1315 (11) |
| MTJ_Sub2 | 1919 (74) | 1224 (64) | 295 (1) |
| MTJ_Sub3 | 1036 (47) | 725 (36) | 311 (11) |
| MTJ_Sub4 | 1502 (62) | 908 (47) | 594 (15) |
| MTJ_Sub5 | 1191 (55) | 793 (48) | 398 (97) |
| MTJ_Sub6 | 824 (29) | 628 (29) | 196 (0) |
| SatCell | 72 (13) | 38 (10) | 34 (3) |
| Fib_Sub1 | 72 (4) | 14 (2) | 58 (2) |
| Fib_Sub2 | 21 (0) | 20 | 1 |
| Fib_Sub3 | 7 (0) | 0 | 7 |
| Fib_Sub4 | 2 (0) | 0 | 2 |
| Fib_Sub5 | 39 (4) | 8 (1) | 31 (3) |
| Fib_Sub6 | 167 (20) | 79 (17) | 88 (3) |
| Fib_Sub7 | 20 (3) | 5 (1) | 15 (2) |
| Fib_Sub8 | 44 (7) | 20 (5) | 24 (2) |
| Fib_Sub9 | - | - | - |
| Fib_Sub10 | - | - | - |
| Smooth_Sub1 | - | - | - |
| Smooth_Sub2 | - | - | - |
| Smooth_Sub3 | - | - | - |
| Endo_Sub1 | 5 (1) | 2 (1) | 3 (0) |
| Endo_Sub2 | 2 (1) | 2 (1) | 0 (0) |
| Endo_Sub3 | - | - | - |
| Endo_Sub4 | - | - | - |
| FatCell | - | - | - |
| Immune_Sub1 | 2 (0) | 0 | 2 |
| Immune_Sub2 | - | - | - |
| Immune_Sub3 | - | - | - |
| Tcell | - | - | - |

To further understand the functional diversity underlying the fibroblast subclusters, we performed GO analysis of the subcluster marker genes. Fib_Sub1 and Fib_Sub3 (MTJ-adjacent) were enriched for actin dynamics, while Fib_Sub2 (fascicular matrix) showed ECM and collagen fibril organization pathways (Data S9). Fib_Sub4 had limited enrichment (Data S9). IFM fibroblasts subtypes displayed both shared and distinct programs: Fib_Sub5 was enriched for cell adhesion and motility; Fib_Sub6 for ECM and collagen organization; Fib_Sub7 for neuronal morphogenesis; and Fib_Sub8 for ECM and vascular development (Data S9). Notably, non-responsive Fib_Sub9 and Fib_Sub10 markers were enriched for neuronal morphogenesis (Data S9). These findings underscore the functional heterogeneity of tendon fibroblasts, particularly among IFM subsets.

### Acute matrisome regulation with exercise at the MTJ primarily occurs in myonuclei

We next examined matrisome gene expression across all cell type subclusters to determine whether ECM remodeling with exercise is a broader, multicellular response. Fibroblasts subclusters collectively regulated a modest 25 matrisome genes with exercise, including upregulation of *PRG4* by the IFM subclusters Fib_Sub6 and Fib_Sub8 and *VCAN* in Fib_Sub6 (Fig. S3 and Data S10). The most transcriptionally responsive subclusters to exercise were the myonuclei subtypes, MyHCI, MyHCII, and MTJ, which together regulated the largest number of matrisome-annotated genes – 235 in total (Table 2 and Data S10). All 14 matrisome elements previously found to be enriched in human hamstring MTJ myonuclei^25^ were regulated by MyHCI, MyHCII and MTJ myonuclei subclusters, including *COL22A1*, which was upregulated by MyHCI_Sub1, MyHCII_Sub2, 3 and 6 and MTJ_Sub1 and 6 (Fig. S8 and Data S10). Satellite cells regulated 13 matrisome genes with exercise, all but *TIMP3* and *CCL2* overlapped with myonuclei matrisome-annotated DEGs. Endothelial subclusters only upregulated *ADAMTS9* and *COL8A1* (Data S10).

To further elucidate the dynamics of the myonuclear matrisome, we performed a comprehensive ECM interactome analysis of all subclusters using MatriCom^37^. Interactions were detected across all cell types in both control and exercised muscle-tendon tissues (Fig. S9A-B and Data S11). However, applying more stringent parameters and restricting the analysis to myonuclei revealed a striking exercise-induced expansion of matrisome interactions, particularly within all MTJ (Sub1-6), MyHCI_Sub1 and MyHCI_Sub5 populations (Fig. 4A-B), consistent with the observed increase in DEGs. Exercise markedly redistributed matrisome activity toward MTJ subclusters, which accounted for ∼72% of all interacting populations compared with ∼52% in controls (Fig. 4C). Most interactions in both conditions were cell-matrisome (Fig. S9C-D). In exercised myonuclei, the baseline repertoire of receptors, integrin α7 (*ITGA7*), β1 (*ITGB1*), and β6 (*ITGB6*), was complemented by *de novo* expression of *CD44* and integrin αV (*ITGAV*) (Fig. 4D). Notably, exercise-challenged MTJ myonuclei acquired a *CD44*+ *CD90*+ (Thy1) surface phenotype and upregulated the transcription factor, *EGR1* (Fig. 4E-F).

**Figure 4.**
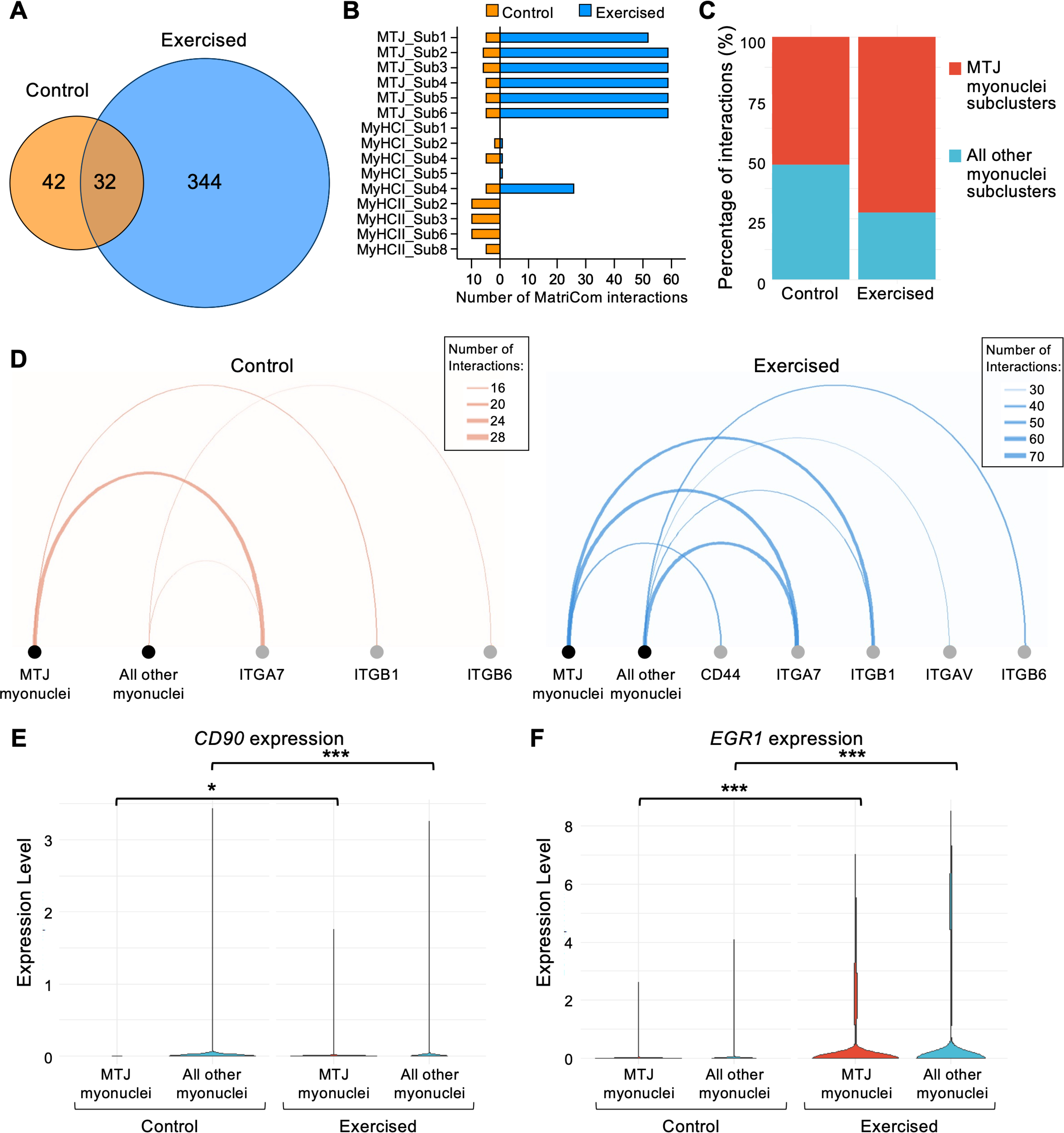
Exercise changes cell-matrisome and matrisome-matrisome communication patterns between myonuclei subclusters. **(A)** Venn diagram showing the number of matrisome interactions (with cells or matrisome), identified using MatriCom^37^, between myonuclei subclusters in control and exercised muscle-tendon tissues. **(B)** Number of MatriCom-identified interactions by myonuclei subclusters in control and exercised samples. **(C)** Percentage contribution of MatriCom-identified interactions from MTJ myonuclei subclusters and all other myonuclei subclusters in control and exercised tissues. **(D)** Chord diagrams showing the number of cell-matrisome interactions via specific cell surface receptors between MTJ myonuclei and all other myonuclei in control and exercised tissues. Line thickness is scaled relative to the total interactions within each condition, not across conditions. **(E-F)** Violin plots showing expression of **(E)** *CD90* (Thy1) and **(F)** *EGR1* in MTJ myonuclei and all other myonuclei. *p=0.033, ***p<0.001 comparing between control and exercised; Wilcox log-rank tests.

## DISCUSSION

Our study represents the most comprehensive single-nucleus resolution dataset of healthy hamstring tendon to date, profiling over 61,000 nuclei from eight human samples, offering greater sample size and tissue coverage than previous single-cell or single-nucleus studies of healthy human tendon^23,25,35,38–41^. Importantly, it also represents the first in vivo investigation of the acute effects of exercise on the human muscle-tendon unit at single-nucleus resolution. The analysis uncovered a robust transcriptional activation in a subset of IFM-resident fibroblasts (Fib_Sub6), implicating them as key mediators of early ECM remodeling within the IFM following mechanical load. In parallel, we identified marked upregulation of MTJ and basement membrane ECM components in MTJ myonuclei, pointing to remodeling of the skeletal muscle connective tissue as a central component of the acute myonuclear response to exercise.

Three mechanosensitive tendon-resident fibroblast populations have been identified, localized adjacent to the MTJ (Fib1: *PIEZO2^+^*, *COL22A1^+^*), in the fascicular matrix (Fib2: *THBSP4^+^*, *COMP^+^*) and in the IFM (Fib3: *PDGFRA^+^*, *VCAN^+^*). Their distinct anatomical locations, and thus differences in nuclear extractability, likely contributed to the observed variation in fibroblast cluster proportions, where Fib1 and Fib3 nuclei from the extended MTJ region isolated via mechanical scraping and the more loosely organized IFM may be more readily released than those embedded sparsely within densely packed collagen fascicles.

Previously, two fibroblast cell types have been identified in human hamstring tendon: *MKX^+^, THBS4^+^, TNMD^+^, PIEZO2^+^* fibroblasts distributed throughout the tissue, and *PDGFRA^+^ NEGR1^+^, NOVA1^+^* fibroblasts enriched in elastin-related genes^23^. Immunostaining showed that the *PDGFRA^+^*fibroblasts were sparse and located near skeletal muscle, however, their origin, tissue specificity, and function remain unresolved, raising questions about whether they represent a tendon-resident population, a muscle-associated cell type or a progenitor-like population shared with other tissues^23^. By integrating marker expression (*VCAN^+^, TNXB^+^, COL14A1^+^, FBN1^+^*), spatial localization to IFM in tissue samples, and their transcriptional response to acute exercise, we further refine their identity as IFM fibroblasts (Fib3). Our dataset also resolved two distinct subsets within the previously described *MKX⁺* fibroblast population. While earlier work identified a single *MKX⁺* subset in human hamstring tendon^23^, our integration of MTJ marker gene expression and spatial localization revealed a MTJ-adjacent *MKX^+^, PIEZO2^+^, THBS4⁺, HHIP^+^, COL22A1⁺* population (Fib1) and a *MKX^+^, THBS4⁺, COMP⁺* fibroblast population within the fascicular matrix (Fib2). The high expression of the mechanosensitive ion channel *PIEZO2*^31^ and expression of MTJ genes^11^ in Fib1 fibroblasts suggest they are uniquely positioned to sense tendon loading. The minimal overlap in modest exercise responses among the three fibroblast clusters further supports their distinct roles in tendon homeostasis.

We also identified the presence of a fourth fibroblast cell type in human hamstring muscle-tendon tissue, characterized by the expression of *TENM2*, *PAK3*, *ITGA6* and *SLC22A3* (Fib4). Despite its relatively high expression, we were unable to confidently resolve the localization of *TENM2* in muscle-tendon tissue sections due to Fib4 constituting to a small portion of fibroblast nuclei. Integration with a skeletal muscle dataset indicated that Fib4 closely resembles perineural fibroblast found in human skeletal muscle^36^. *TENM2+* fibroblasts have also been identified in single-cell studies of mouse skeletal muscle^42,43^. Together with the GO enrichment analysis of Fib4 marker genes, these data support Fib4 as being a possible skeletal muscle-resident, perineural fibroblast.

IFM subclusters (Fib_Sub5-8) exhibited diverse transcriptional programs in response to exercise. Together with GO analysis of subset marker genes, our findings highlight greater functional heterogeneity within the IFM compartment. Consistent with this, a previous single-cell study of young and old equine tendons reported transcriptional heterogeneity among *PRG4⁺* IFM fibroblasts, identifying three IFM clusters and two *THBS4⁺, COMP⁺* fascicular clusters ^22^. Notably, both the prior study and the present work converged on an ECM-enriched IFM fibroblast population: *“IFM A*” in horse tendon and Fib_Sub6 in our dataset^22^. Extending this observation, our data reveal that Fib_Sub6 mounts the most robust transcriptional response to exercise, with upregulation as well as downregulation of genes, suggesting this subset may play a central role in early ECM remodeling within the IFM following mechanical load. Interestingly, the *“IFM A”* population was the most affected with age^22^. We attempted to integrate our dataset with the published equine tendon single-cell data to determine whether *“IFM A”* and Fib_Sub6 represent analogous fibroblast populations across species. However, differences in nuclei versus whole cells and/or species differences prevented successful alignment of the datasets.

In contrast to the IFM, the fascicular matrix fibroblast subcluster enriched in genes regulating ECM organization (*COMP^+^* Fib_Sub2) was the least transcriptionally responsive to exercise. This aligns with the distinct biological dynamics of the two tendon compartments, where the glycoprotein-rich ECM of the IFM exhibits higher turnover than the collagen-rich fascicular matrix^14,44^, which was also demonstrated by the exercise-induced upregulation of proteoglycans, *PRG4* (lubricin) and *VCAN* (versican) in Fib_Sub6. Fascicular matrix fibroblasts (Fib_Sub1-3) showed a consistent downregulation of co-chaperones in response to exercise. Among these were C*HORDC1* (cysteine- and histidine-rich domain-containing protein 1) that interacts with the chaperone Hsp90 (HspC) in high ADP:ATP ratios^45^, *DNAJB6* (encoding Hsp40 member 6) a co-chaperone of Hsp70, best known for preventing protein aggregation in neurons and muscle fibers^46,47^, and *PTGES3* (p23), a co-chaperone of Hsp90^48^. These molecular chaperones are essential for folding of new and misfolded proteins and protein trafficking, which can be disrupted because of environmental stress, such as temperature and mechanical loading. While their specific roles in tendon fibroblast responses remain unclear, the observed transcriptional downregulation suggests a homeostatic, protective adaptation rather than an anabolic one. This supports previous observations that tendons require prolonged or repeated loading stimuli to drive net increases in collagen production^49^. However, some of the observed transcriptional changes, such as the differential expression of *HNRNPU*, a key factor in X chromosome inactivation via the long noncoding RNA, *XIST*^50^, reflect differences in sex distribution between groups rather than direct effects of exercise. This highlights an important limitation of the study, the unequal representation of sexes in the exercised versus control groups, which may confound interpretation of sex-linked or sex-biased gene expression. While we aimed for equal sex distribution and deliberately included female donor samples, technical issues such as sample quality and post-sequencing QC thresholds resulted in a skewed sex ratio in the final dataset.

By focusing on early time points following a single bout of exercise, our study was designed to capture the immediate transcriptional responses and identify mechanosensitive tendon cells. While this approach effectively reveals acute regulatory pathways, longer-term adaptation mechanisms remain to be explored. Future studies incorporating extended time courses or repeated loading regimens will be important to fully understand the extent of remodeling in tendon adaptation.

Our study provides the first evidence that myonuclei at the MTJ mount a robust transcriptional response to a single bout of heavy eccentric resistance exercise. Although not the primary target of this loading modality, slow-twitch type I myonuclei still responded markedly, with 1365 DEGs in one MyHCI subset (Table 2). For comparison, the most responsive fibroblast subcluster (Fib_Sub6) exhibited 167 DEGs. Due to the sample preparation method, which involved scraping away the muscle fibers, all myonuclei profiled in this study originated from the MTJ. A previous study reported that 26% of type I and 19% of type II MTJ myonuclei express *COL22A1*^9^. In line with this, ∼26% ± 7% (mean ± SD; n=8) of myonuclei in the present study expressed *COL22A1*. The remaining myonuclei clusters expressed *NCAM1*, which was previously found to be restricted to MTJ and transitional MTJ myonuclei subclusters^25^, indicating that the majority of our myonuclei are derived from the transitional MTJ region, as opposed to myonuclei from the muscle belly. The role of the transitional MTJ zone in MTJ maintenance and resilience to injury remains unknown, so our data provide valuable insight into the acute response of this zone to eccentric exercise. The acute and robust upregulation of *COL22A1* and *LAMA2* in myonuclei highlights basement membrane remodeling as a central component of early MTJ adaptation to heavy eccentric exercise. Furthermore, the rapid upregulation of *MYOM3* and *RBM20*, a potent regulator of titin splicing^51^, indicates sarcomere remodeling at the myofiber tips. Similarly, the 38 upregulated and 34 downregulated genes in satellite cells at the MTJ provide valuable insight into the early myogenic response in humans in vivo.

MatriCom analysis^37^ enabled a global, unbiased inference of ECM network remodelling with exercise, integrating all detectable ECM-ECM and cell-ECM interactions across cell populations. De novo expression of *CD44* and integrin αV in MTJ myonuclei with acute exercise is suggestive of ECM remodeling at myofiber tips in contact with the tendon and consistent with recent descriptions of tendon repair-associated molecular programs seen in mice^52,53^. Notably, MTJ myonuclei also upregulated *CD90* and *EGR1*, hallmarks of a reactive state in which homeostatic and mechanosensitive pathways converge to preserve tendon integrity^52^. In agreement with this shift, the MatriCom analysis also revealed increased expression of *VEGFA*, the basement membrane proteoglycan perlecan (*HSPG2*) and the integrin-binding nephronectin (*NPNT*) in MTJ myonuclei with acute exercise (Data S11). How these proteins contribute to MTJ and tendon organization remains undefined.

Interestingly, although not the primary focus of this study, we detected transcriptional changes in a few circadian clock genes (e.g., *ARNTL*, *CRY1*) in subsets of IFM (Fib_Sub6 and Fib_Sub8) and MTJ-adjacent (Fib_Sub1) fibroblasts following acute exercise (Data S3). This is consistent with previous work showing that mechanical cues can act as zeitgebers in peripheral tissues, especially in skeletal muscle^54^, also demonstrated by regulation of multiple clock genes in all exercise-responsive myonuclei in our dataset (Data S3). Although further studies are needed to elucidate the functional relevance of these changes in tendon, the consistent timing of the intervention and tissue collection adds confidence to our observation of exercise-induced shifts in clock gene expression. Together, our findings support a potential link between mechanical loading and circadian regulation in tendon, a relationship that may influence collagen homeostasis^55,56^.

This study provides a foundational single-nucleus atlas of the human muscle–tendon unit’s acute response to mechanical load, revealing spatially distinct ECM programs in fibroblasts and myonuclei. The coordinated yet cell type-specific responses highlight dynamic differences between ECM niches of the MTJ and tendon, offering a basis for investigating long-term adaptation and remodeling in these compartments.

## ACKNOWLEDGEMENTS

Fluorescence-activated nuclei sorting was performed at the University of Copenhagen Biotech Research & Innovation Centre FACS Facility with help from Dr. Rajesh Somasundaram. Single-nuclei libraries were prepared at the University of Copenhagen Biotech Research & Innovation Centre Single-Cell Core Facility. Microscopy was performed at the University of Copenhagen Core Facility for Integrated Microscopy. The authors would like to thank the Bispebjerg Hospital Department of Orthopedic Surgery, Ann-Christina R Reimann, Dr. Grith Højfeldt, Christian Hoegsbjerg, Stine Bitsch-Olsen, Dr. Costanza Montagna and Max FR Merkel from the Institute of Sports Medicine Copenhagen for their generous assistance with this study. We would also like to thank Dr. Chavaunne Thorpe (Royal Veterinary College, London) for valuable discussions on interfascicular matrix cell biology, enabled through a Tendon Regeneration Network (TENET) European Cooperation in Science and Technology Action Short-Term Scientific Mission.

## FUNDING

This work was supported by a Lundbeck Foundation Grant (R198–2015-207 to M.K.), a Novo Nordisk Foundation Project Grant (NNF23OC0086015 to C.-Y.C.Y.) and an Agility Project Award from the Wu Tsai Human Performance Alliance and the Joe and Clara Tsai Foundation (2023 to M.K. and C.-Y.C.Y.). K.K. and M.Y.B. were supported by the Novo Nordisk Foundation Hallas-Møller Investigator grant (NNF21OC0067146 to K.K.).

## AUTHOR CONTRIBUTIONS

C.-Y.C.Y., M.K., A.L.M. and M.R.K. conceived and designed the study. C.-Y.C.Y. supervised the study. A.M., D.S. and C.-Y.C.Y. performed the experiments. P.S. designed the snRNAseq data quality control and analysis workflow. V.I. performed the MatriCom analysis. A.M., D.S., P.S., V.I. and C.-Y.C.Y. formally analyzed and curated the data. J.R.J. recruited, supervised the training of subjects and collected the tissue samples. C.-Y.C.Y., A.J.U., M.Y.B. and K.K. participated in the single nucleus isolation and RNA library preparation. C.-Y.C.Y. drafted the manuscript. All authors reviewed and approved the manuscript.

## DECLARATION OF INTERESTS

The authors declare no competing interests.

## DATA AVAILABILITY

The snRNAseq data are deposited in the ArrayExpress database with accession numbers E-MTAB-15400. All other relevant data can be found within the article and its supplementary information.

## MATERIALS AND METHODS

### Experimental Design

The primary objective was to determine the response of fibroblasts in the human muscle-tendon unit to acute heavy resistance exercise. The secondary objective was to determine the localization of these cell types. Participants undergoing anterior cruciate ligament (ACL) reconstruction surgery were recruited to perform a single session of eccentric hamstring exercises or no exercise (controls), and their hamstring tendons were harvested 4 h later. While the initial recruitment target was six biological replicates per group, quality control filtering resulted in a final dataset comprising four biological replicates per group for snRNAseq. In situ hybridization by RNAScope was performed in three biological replicates per group. No power calculations were performed, as this was an exploratory study.

### Ethics

Written and informed consent was obtained from all participants (ethics approval H-18022988) by the Regional Ethical Committee for the Hospital Region of Greater Copenhagen, in accordance with the Declaration of Helsinki II. Additional control samples were collected for RNAscope experiments (ethics approval H-20044907). The study was reported to the Danish register (Datatilsynet) and was performed in accordance with Danish law (Lov om behandling af personoplysninger).

### Human participants

Patients with a scheduled ACL reconstruction surgery were invited to participate in the study. Only those assigned to the second surgical slot of the day (starting 11:00) were considered for inclusion to allow precise timing of the pre-operative intervention. Participation was restricted to individuals who had not engaged in regular hamstring-targeted resistance exercise within the preceding three months. Twelve patients (5 females and 7 males) were enrolled and randomized into either a training group or a control group (see Table 1). The control group did not perform any exercise on the day of the surgery, and the training group performed one single session of eccentric exercise (see below for details) 4 hours before the scheduled surgery. One sample from the control group and one sample from the exercise group did not contain sufficient nuclei for snRNAseq, leaving 5 samples in the control group and 5 samples in the exercise group to be processed for snRNAseq. After initial quality control analysis of the sequenced samples, one sample in the control group contained mostly doublets and one sample in the exercise group showed problems with the cDNA library generation and so were excluded from the final data analysis. See Table 1 for age and sex distribution of the final samples analyzed (n=4).

### Resistance exercise

A resistance training protocol targeting the hamstring muscles (semitendinosus and gracilis), with an emphasis on heavy eccentric loading, incorporating unilateral and bilateral exercises performed to fatigue under supervised conditions, as previously described^12^. In brief, starting ∼7:00, patients warmed up for 5 min on an exercise bike and then performed 3 sets of 8 repetitions of the following exercises until exhaustion, unilateral eccentric leg curl (on the operation leg only) in the prone position to exhaustion, bilateral leg press, bilateral Nordic hamstring leg curl, bilateral leg extension, unilateral eccentric leg curl (on the operation leg only) in the prone position to exhaustion, and stiff-legged deadlift performed to exhaustion during the lowering phase. Participants rested for two minutes between the different exercises. All exercises were performed under supervision to ensure a proper form and steady tempo was maintained, and all patients were supervised by the same trainer.

### Human tissues

Healthy muscle-tendon tissues (gracilis and semitendinosus) were obtained as waste tissue from ACL reconstruction surgeries (see Table 1). Within 8 minutes of excision (∼11:15), the tissue was immediately placed in ice-cold phosphate-buffered saline (PBS) and kept on ice. To minimize the presence of myonuclei in our dataset, as our focus was on the maintenance of fibrous connective tissue, all visible attached muscle tissue was carefully dissected from the tendon by scraping with a scalpel. This procedure was performed on ice to minimize RNA metabolism. Our pilot experiment confirmed that this method preserved the abundant myonuclei localized to the MTJ. Following dissection, which took ∼30 min, tendon tissue (100-500 mg) was cut into ∼50 mg pieces and snap-frozen in individual cryotubes with liquid nitrogen. For RNAscope, muscle-tendon samples were embedded, without scraping, in Tissue-Tek (Sakura Finetek) and frozen in liquid nitrogen-cooled isopentane. All tissues were stored at −80°C until further processing.

### Preparation of single nuclei suspension

An adapted version of the ‘Frankenstein’ protocol^57^ was used to prepare nuclei suspensions from frozen tendon tissue samples^58^. In brief, snap frozen tendons were cut into smaller pieces (∼0.5 mm^3^) in a −20°C cabinet and then were transferred to 2-ml screw cap tubes, with each tube containing ≤50 mg tissue and five 2.3 mm-diameter stainless steel ball bearings (BioSpec Products Inc., Bartlesville, OK) (Table 1). Samples were kept on dry ice to prevent thawing. All further steps were carried out on ice or at 4°C using chilled reagents. After, 0.5 ml Nuclei EZ Lysis Buffer (Sigma) was added to each tube and then homogenized using a FastPrep-24 (MP Biomedicals) at 4.0 m/s for 20 s, cooled down on ice for 5 min, then homogenized once more at 4.0 m/s for 20 s and cooled down on ice for 5 min. The homogenate was transferred to a pre-chilled 2-ml microcentrifuge tube, 0.5 ml chilled Nuclei EZ Lysis Buffer was added to each tube, mixed gently using a wide bore pipette, and incubated on ice for 5 min with two further mixes using a wide bore pipette during the incubation. The homogenate was then filtered through a 70 µm mini strainer (PluriSelect) and centrifuged for 5 min at 500 x g at 4°C. After, the pellet was resuspended in 1.5 ml chilled Nuclei EZ Lysis Buffer by pipetting 10 times and centrifuged for 5 min at 500 x g at 4°C. After, the pellet was equilibrated on ice with 0.5 ml nuclei wash and suspension buffer (2% BSA in PBS (Sigma), 2 mM MgCl_2_ (Sigma), 0.2 U/µl Protector RNase inhibitor (Roche)) for 5 min. After, 1 ml nuclei wash and suspension buffer was added, the pellet was then resuspended by pipetting 20 times using a 1000 µl pipette tip and then centrifuged for 5 min at 500 x g at 4°C. The pellet was resuspended in 1 ml nuclei wash and suspension buffer by pipetting 20 times using a 1000 µl pipette tip, then filtered through a 40 µm mini strainer (PluriSelect). The filter was washed with 0.5 ml nuclei wash and suspension buffer. For each sample, the suspension was split into two 1.5 ml microcentrifuge tubes containing 20% and 80% of the volume, respectively. The tubes were centrifuged for 5 min at 500 x g at 4°C and the pellets were each resuspended in 200 µl nuclei wash and suspension buffer. Trypan blue (Sigma) and a hemocytometer were used to count the number of nuclei in 5 µl of the suspension containing 20% nuclei. Nuclei suspensions were stored on ice for no more than 30 min before nuclei sorting.

### Fluorescence-activated nuclei sorting

Nuclei were sorted by fluorescence-activated cell sorting (FACS) to remove non-nuclear materials and any nuclear aggregates. To the tube containing 80% nuclei suspension, 1.5 µg/ml 7-aminoactinomycin D (7AAD), a fluorescent intercalator that associates with DNA, was added. The tube containing 20% nuclei suspension was used as a negative label control to set up sorting gates. Forward scatter/side scatter gates were used to remove clumps of cells and debris. Single 7AAD-labelled nuclei were sorted into a 5°C-chilled 1.5 ml microfuge tube coated with nuclei wash and suspension buffer. FACS was performed on a FACSAria III (BD Biosciences). Sorted nuclei suspensions were transported on ice and immediately used for single nuclei cDNA library preparation. Five samples in the control group and five samples in the exercise group were processed for snRNAseq.

### 10X Chromium and next generation sequencing

The official Chromium Single Cell 3’ Reagents Kits v3 User Guide (version CG000183 Rev A) was followed for the creation of single nuclei cDNA libraries. Target nuclei recovery was 10,000 per sample. Sequencing of the ten libraries was performed using Illumina HiSeq by Genewiz GmbH (Leipzig, Germany). The samples were sequenced in three batches: first, one pilot from the control group, second, one sample from each group, and third, the remaining samples were sequenced after an additional step of primer dimer removal. The final samples analyzed were from 4 biological replicates in each group.

### snRNAseq data analysis

Sequencing files were demultiplexed, barcodes processed, mapped to the human genome (Gencode release 38 (GRCh38.p13))^59^, and reads counted using Cell Ranger v7.0.0. Two samples were excluded from further analysis because of lower mapping frequency and higher than expected nuclei count. The presence of ambient RNA was identified by examining the frequency distribution of total UMI counts and partially corrected for using CellBender (0.2.0)^60^. The ambient corrected RNA counts were used to test for high levels of ribosomal or mitochondrial RNA, which should not be present in snRNA data. Scrublet (version Dec2020 and python version 3.6)^61^, which calculates a doublet score based on the observed and synthetic transcriptomes in PCA space, was run three times on each data set to identify and remove doublets. The final data set contained 3550-11879 nuclei per sample. Seurat (version 4.1.1)^62^ was used for analysis of the cleaned data sets. Data sets were normalized with SCTransform and integrated using Seurat Anchors (v3). For the main analysis, clustering was performed using 30 PCs on integrated data at a resolution of 0.3. This resulted in 18 clusters which separated clusters on the UMAP without splitting sensible clusters into fractions. The clusters were then given names based on expression of known marker genes. To fine tune the clustering, the clusters that belonged to a broader visual cluster (e.g., different myonuclei clusters) were grouped together and re-clustered at a resolution of 0.3. Cluster markers were determined with the function FindConservedMarkers. The expression of each gene was compared to other clusters and only significant markers (log2 fold change >1, FDR <0.05) present in each sample were accepted. For the subcluster search, two different comparisons were made with (1) one cluster versus all other cluster using FindConservedMarkers, and (2) one cluster against only other subclusters in the same subgroup using the less stringent FindMarkers. The clusters were compared to clusters in other snRNAseq data sets using the function FindTransferAnchors (“SCT”, reduction = “pcaproject”, reference.reduction = “pca”).

GO analysis for biological pathways in subclusters (Data S4 and Data S9) was performed using topGO(ref) (options: classic, fisher)^63^. Core matrisome and matrisome-associated genes were identified using the human matrisome list defined by Naba *et al.*, 2012 (Hs_Matrisome_Masterlist_ Naba et al_2012)^64^, available from The Matrisome Project. Protein-protein interaction networks and GO analysis for biological pathways in fibroblast clusters (Data S7) were generated using the STRING database (version 12.0)^65^. Only interactions with a medium confidence score (≥ 0.4) or higher were included, and both known and predicted interactions were considered. MatriCom analyses were run with mostly default settings^37^ on a local installation of the application, barring the percentile of cells expressing interaction partners, which was set at 80% to increase stringency, and reliability levels were set to all possible in the exploratory phase to increase the likelihood of discovering matrisome interactions. For the myonuclei-focused analysis, reliability was set at level 3, the maximum, to further support stringency.

### Exercise effect on cell types and transcripts within cell types

To determine whether exercise influenced the number of nuclei that belonged to each cluster, the exercise data was pseudobulked (aggregated by subject) within each cluster. DESeq2^66^ was used to make statistical comparison on nuclei counts per cluster (versus non-pseudobulked marker analysis on gene expression per cluster). The criteria for DEG were set with alpha = 0.05 and log2 fold change = 0.585. Similarly, DESeq2 was used to compare the number of bulk transcript counts that belong to each cluster and test whether they are up or downregulated by exercise.

### Histology

Longitudinal sections of tendon were stained with hematoxylin and eosin. Fresh frozen tissues of 12-µm thickness were air dried, then stained with Mayer’s hematoxylin for 1 min, washed with tap water for 5 min, then stained with eosin for 5 min, and washed briefly in tap water. Sections were then dehydrated through an ethanol series and xylene before mounting in DPX mounting medium (Merck). Images were captured on an Olympus BX51 microscope (Olympus, Japan) with CellSens software, using a 10× objective.

### RNAscope in situ hybridization

RNA in situ hybridization was performed using RNAscope Multiplex Fluorescent Reagent Kit v2 and RNAscope reagents (ACD, supplied by Bio-Techne Ltd, UK, unless specified otherwise) following the manufacturer’s protocol for fresh frozen tissues. Tissues from seven biological samples were used: 3 females and 4 males, 30 ± 8 years (mean ± SD; range 24-46). Donor age and sex information for each RNAscope experiment are provided in Data S12. Longitudinal tissue sections of 8-µm thickness were fixed with 10% neutral buffered formalin for 1 h, at 4°C, then dehydrated through an ethanol series (50%, 70%, 100% twice, for 5 min each at RT) and stored in 100% ethanol at −20°C overnight. After, the slides were dried for 5 min and outlined with a hydrophobic barrier pen, before addition of hydrogen peroxide for 10 min, followed by washing in distilled water. Tissues were permeabilized with Custom Pretreatment Reagent for Bone (300040) for 30 min at 40°C, followed by two washes with PBS before probe hybridization. The RNAscope tissue pretreatment protocol was optimized for tendon tissue and effectively removed cytoplasmic proteins from muscle, as indicated by loss of eosin staining, but myonuclei remained intact and detectable (Fig. S4A-B). Detection probes against *COL22A1* (818811-C3), *COMP* (457081-C2), *MYH1* (1242761-C3), *MYH2* (504731-C1), *MYH7* (508201-C2), *PDGFRA* (604481-C2), *TENM2* (1556761-C1), and *THBS4* (426591-C1), which detect both pre- and mature RNAs, were applied to the tissue sections for 2 h at 40°C and then stored overnight at room temperature in 5× saline-sodium citrate buffer (0.75 M socdium chloride, 75 mM trisodium citrate, pH 7.0). The following day, the signal amplification was performed according to the manufacturer’s protocol. Incubations were carried out at 40°C, followed by two 2 min washes with RNAscope wash buffer. Amplifier incubations were 30 min and 15 min for the two first and the third amplifiers, respectively. Horseradish peroxidase incubation for 15 min was followed by 30 min incubation with a tyramide dye fluorophore (OPAL 690, FP1497001KT, Akoya Biosciences, Marlborough, USA) diluted 1:1000 in RNAscope TSA dilution buffer and 30 min incubation with a horseradish peroxidase blocker. The horeseradish peroxidase, fluorophore, and blocking steps were repeated with another tyramide dye fluorophore (OPAL 570, FP1488001KT, Akoya Biosciences, Marlborough, USA). Probes labelled with OPAL 690 were pseudocolored red and probes labelled with OPAL 570 were pseudocolored yellow in the images. Nuclei were labeled with RNAscope 4′,6-diamidino-2-phenylindole (DAPI) for 30 seconds directly after the final wash and slides were mounted with ProLong Gold Antifade mounting medium (Invitrogen) and a cover glass. Images were captured by confocal laser scanning microscopy.

### Confocal laser scanning microscopy

Images of RNAscope-stained longitudinal sections were captured using laser scanning confocal microscopy with a Zeiss LSM780 microscope (Zeiss, Jena, Germany) at a resolution of 0.12 µm per pixel with an optical thickness of 0.6-0.8 µm (pinhole 1 AU). An LD Plan-Neofluar 20x (0.8 NA) objective and Zeiss Zen Black 2012 software were utilized. A 405nm diode (30mW), 543nm HeNe (1.2mW), and 633nm HeNe (5mW) lasers were used to excite DAPI, Opal 570, and Opal 690, respectively. Four times averaging was used to reduce noise during image acquisition.

### RNAscope staining quantification

Quantification was performed in FIJI (ImageJ version 1.54p). Maximum intensity projections were generated from 8 µm z-stacks, 1-µm apart, reducing dimensionality while preserving signal intensity. The interfascicular matrix (IFM) was manually drawn in each image using the drop in autofluorescence of the tissue as a guide (Fig. S4C). Nuclear regions-of-interests (ROIs) were created using the DAPI signal and the “Analyze Particles” function. These ROIs were subsequently divided into an inside and outside IFM group based on spatial overlap with the IFM ROI. RNAscope signals were thresholded using either the Triangle or default algorithm, applied per image to account for background variation. The resulting binary mask was used to determine nuclear gene expression. A nucleus was considered positive if a defined percentage of its area overlapped with the thresholded signal. Based on staining characteristics of each probe, the threshold was set at 5% for *THBS4* and *COMP* (high expression, nuclear and cytoplasmic), and 0.25% for *COL22A1* and *PDGFRA* (small punctate nuclear staining). Mixed-effect logistic regression was used (performed in R with lme4^67^) to test for significant differences in RNAscope experiments. This approach models the probability of a nucleus being positive for a given transcript (binary outcome) while accounting for the total number of nuclei analysed and for inter-sample variability through inclusion of sample identity as a random effect. This framework is robust to sparse positive events, as it leverages the full denominator of nuclei screened rather than comparing only raw counts.

### Statistical analyses

Statistical analyses of snRNAseq data are outlined in the corresponding methods sections. Sample sizes, statistical tests used, significant differences, and p or FDR values are indicated in the figures or noted in the corresponding figure legends.

## Notes

### Competing Interest Statement

The authors have declared no competing interest.

