## Supplementary material for "Spatially distinct ECM-producing fibroblasts and myonuclei orchestrate early adaptation to mechanical loading in the human muscle-tendon unit": Fig. S1-9

**List of Supplementary Files for:**

#### SUPPLEMENTARY FIGURES

Figure S1. Sample and condition distribution across clusters. (PDF)

Figure S2. Myosin expression by myonuclei clusters and subclusters. (PDF)

Figure S3. Cluster-enriched matrisome genes in tendon fibroblast clusters. (PDF)

Figure S4. RNAscope localization of transcripts in tendon sections. (PDF)

Figure S5. Cross-dataset integration and correlation of fibroblast clusters. (PDF)

Figure S6. Exercise-induced differential regulated RNAs in fibroblast clusters. (PDF)

Figure S7. Transcriptomic characterization of hamstring tendon fibroblast subclusters. (PDF)

Figure S8. Exercise-induced matrisome changes in myonuclear subtypes and their overlap with MTJ-enriched genes. (PDF)

Figure S9. Matrisome communication between nuclei subclusters in human hamstring muscle-tendon tissue. (PDF)

#### SUPPLEMENTARY DATA FILES

Data S1. Lists of marker genes of all main clusters. (.XLSX)

Data S2. Lists of marker genes of all subclusters. (.XLSX)

Data S3. Lists of exercise-induced DEGs in subclusters. (.XLSX)

Data S4. Gene ontology biological pathways analysis of exercise-induced DEGs of subclusters. (.XLSX)

Data S5. List of marker genes from within main cluster comparisons. (.XLSX)

Data S6. Lists of exercise-induced DEGs in main clusters. (.XLSX)

Data S7. Gene ontology biological pathways from STRING analysis of exercise-induced DEGs in fibroblast clusters. (.XLSX)

Data S8. Lists of subcluster marker genes from within subcluster comparisons. (.XLSX)

Data S9. Gene ontology biological pathways analysis of subcluster marker genes (.XLSX)

Data S10. Lists of matrisome-annotated exercise-induced DEGs in all subclusters. (.XLSX)

Data S11. List of cell-matrisome and matrisome-matrisome interactions between all subclusters. (.XLSX)

Data S12. Source data for generating the figures. (.XLSX)

### Møbjerg *et al.* – Supplementary Figure 1

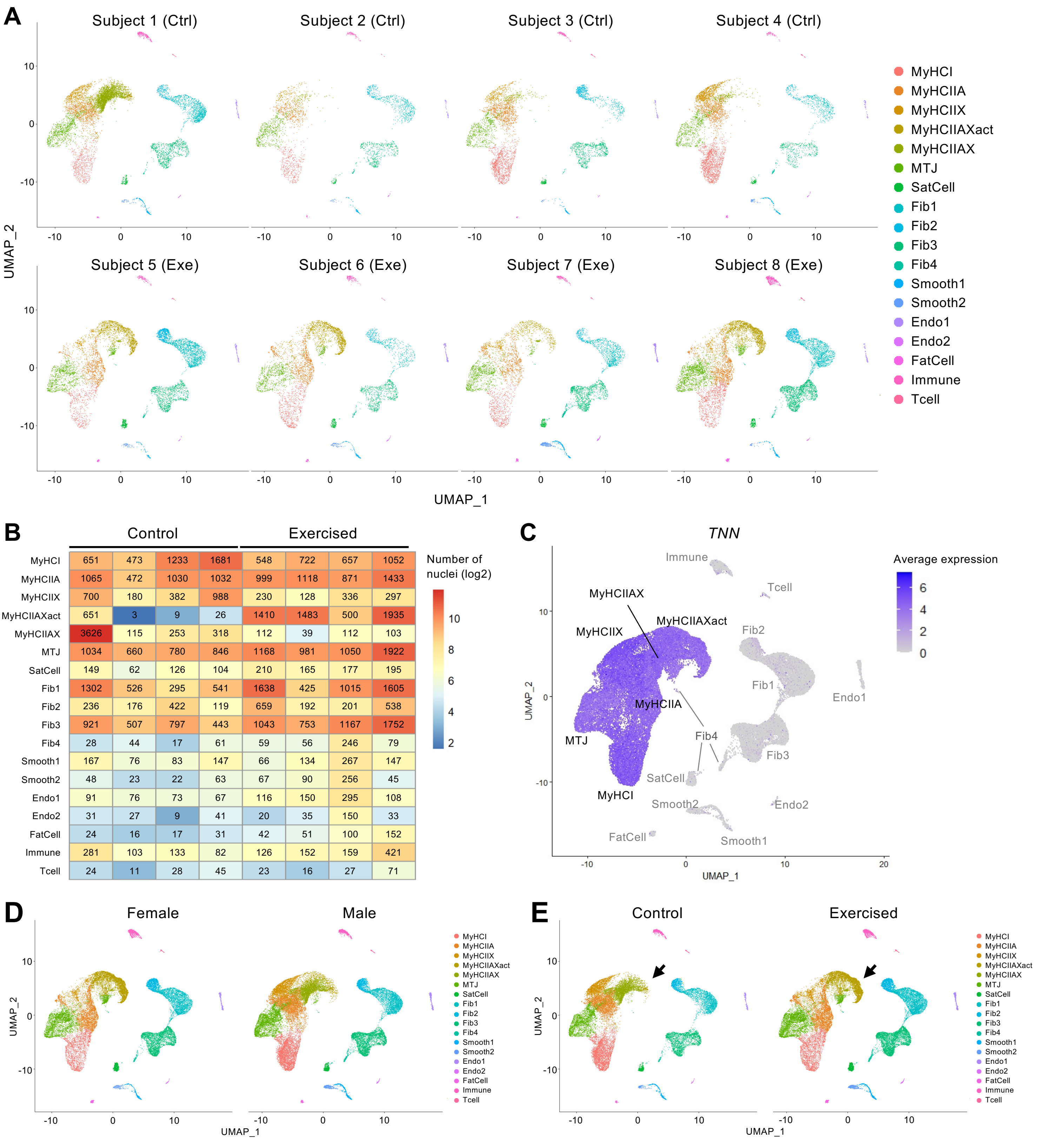


**Figure S1. Sample and condition distribution across clusters.** **(A)** UMAP of the 18 clusters in each of the eight samples analyzed (4 controls (Ctrl) and 4 exercised (Exe)). **(B)** Heatmap of the nuclei distribution across the eight samples. Color intensity represents log2-transformed values, while absolute nuclei counts are displayed within each box. **(C)** UMAP feature plot of *TTN* expression restricted to myonuclei in all samples (n=8). **(D)** UMAP of the 18 clusters separated by sex (3 females, 5 males) and **(E)** by mechanical loading (4 controls, 4 exercised). Arrows point to the MyHCIIAXact cluster, which increased with exercise.

### Møbjerg *et al.* – Supplementary Figure 2

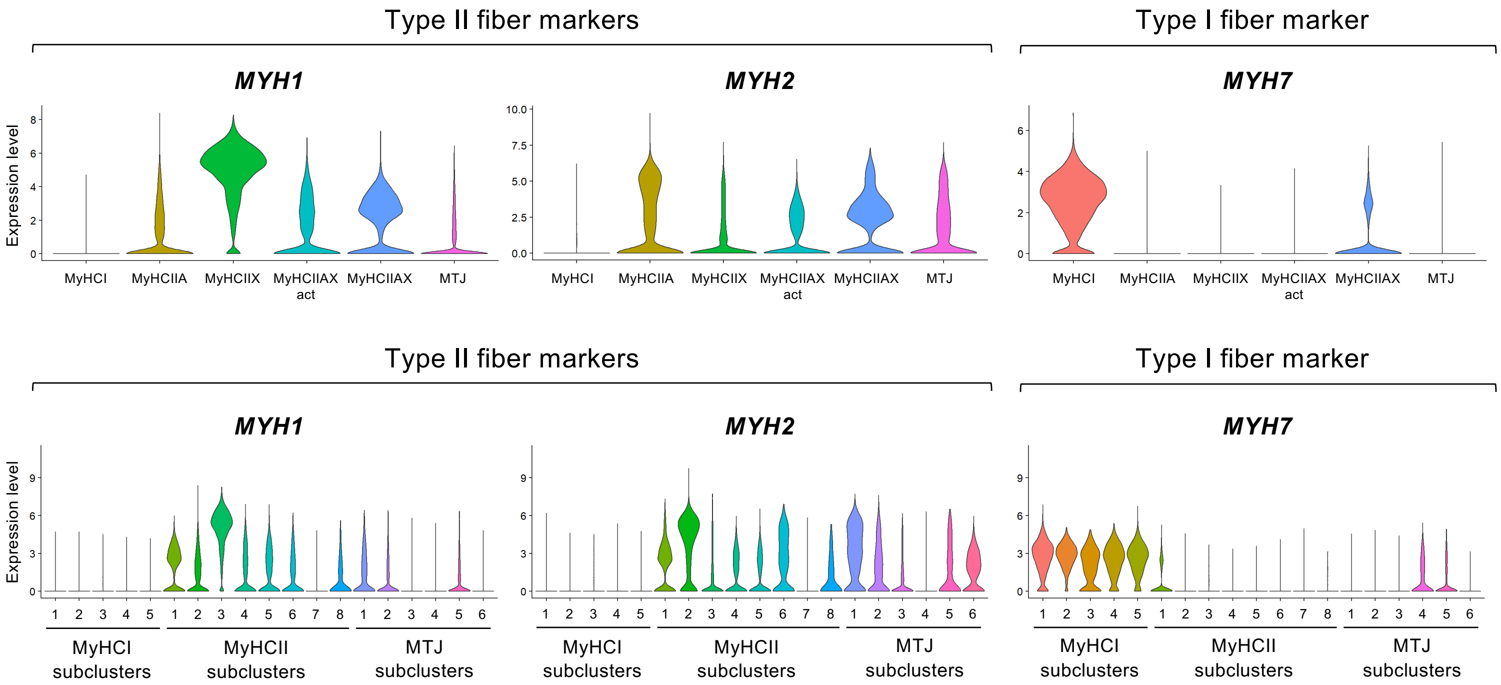


**Figure S2. Myosin expression by myonuclei clusters and subclusters.** Violin plots showing *MYH1*, *MYH2* and *MYH7* expression that distinguishes between type I (*MYH7*) and type II (*MYH1* and *MYH2*) fibers.

### Møbjerg *et al.* – Supplementary Figure 3

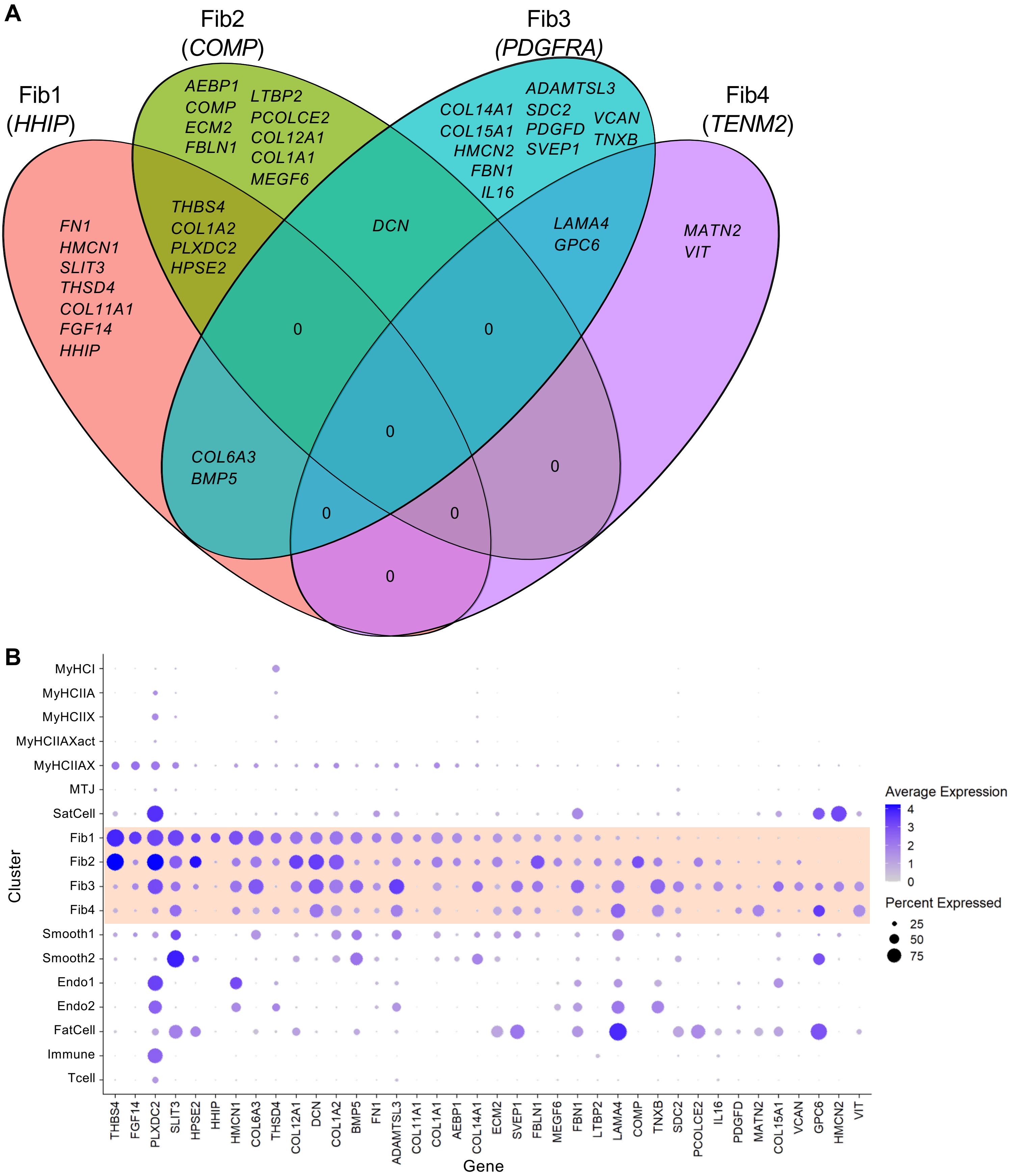


**Figure S3. Cluster-enriched matrisome genes in tendon fibroblast clusters. (A)** Venn diagram illustrating the minimal overlap of core matrisome and matrisome-associated markers across the four fibroblast nuclei clusters. **(B)** Dot plot of the 38 matrisome-annotated genes expressed by fibroblast clusters (highlighted in orange). Expression values displayed are derived from RNA assay data obtained from individual nuclei.

### Møbjerg *et al.* – Supplementary Figure 4

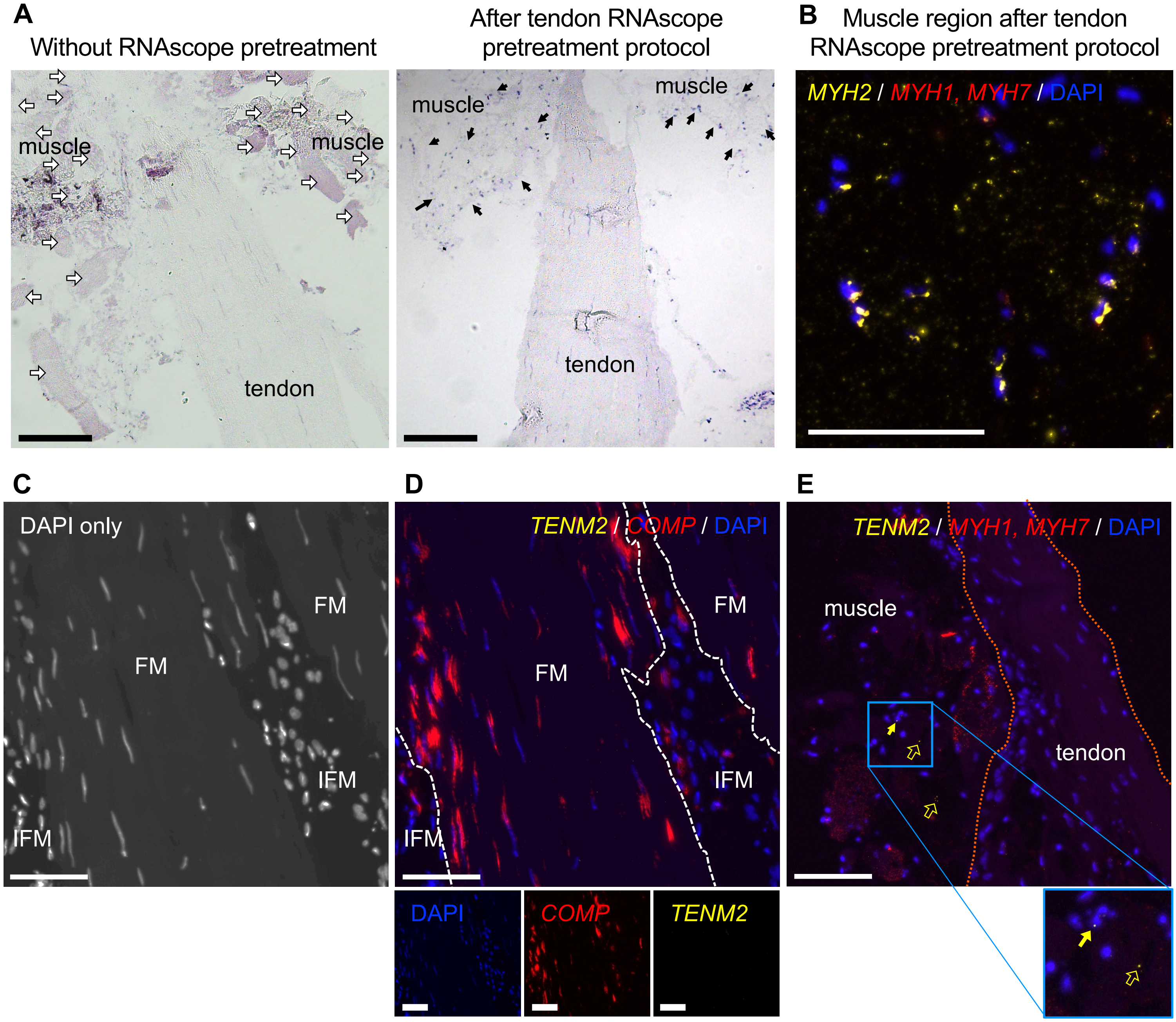


**Figure S4. RNAscope localization of transcripts in tendon sections. (A)** Example of H&E-stained tendon sections, without and with RNAscope pretreatment, used to identify tissue-type (muscle, tendon or MTJ) and guide RNAscope staining in subsequent serial sections. After tissue permeabilization, the tendon fascicles, including MTJ-adjacent areas, are intact. White arrows point to muscle fibers, visible without RNAscope pretreatment. Black arrows point to regions where myonuclei that remain intact after tissue permeabilization. **(B)** Example image confirming the detection of human skeletal muscle-specific RNAs, *MYH1* (red), *MYH2* (yellow) and *MYH7* (red) in myonuclei of tissue sections after the tendon RNAscope pretreatment. **(C)** Example image showing the background fluorescence from the DAPI signal from the tendon fascicles being used to guide the creation of IFM regions-of-interest (ROIs), marked by the dotted white line in (D). **(D)** Representative image of a longitudinal section of human hamstring tendon tissue stained with RNAscope probes to human *COMP* (red) and *TENM2* (yellow) and DAPI to stain nuclei. No positive staining for *TENM2* was observed in tendon. **(E)** Representative image of a longitudinal section of human hamstring muscle-tendon tissue stained with RNAscope probes to human *MYH1* and *MYH7* (both red) and *TENM2* (yellow) and DAPI to stain nuclei. No consistent positive staining for TENM2 was observed except for one instance where signal was detected in the muscle region of the tissues (closed yellow arrow for nuclei-associated signal and open yellow arrows for non-nuclei-associated). N=3 biological replicates stained. FM, fascicular matrix. IFM, fascicular matrix. Orange dotted lines indicate the borders of the tendon tissue. Scale bars, 100 µm.

### Møbjerg *et al.* – Supplementary Figure 5

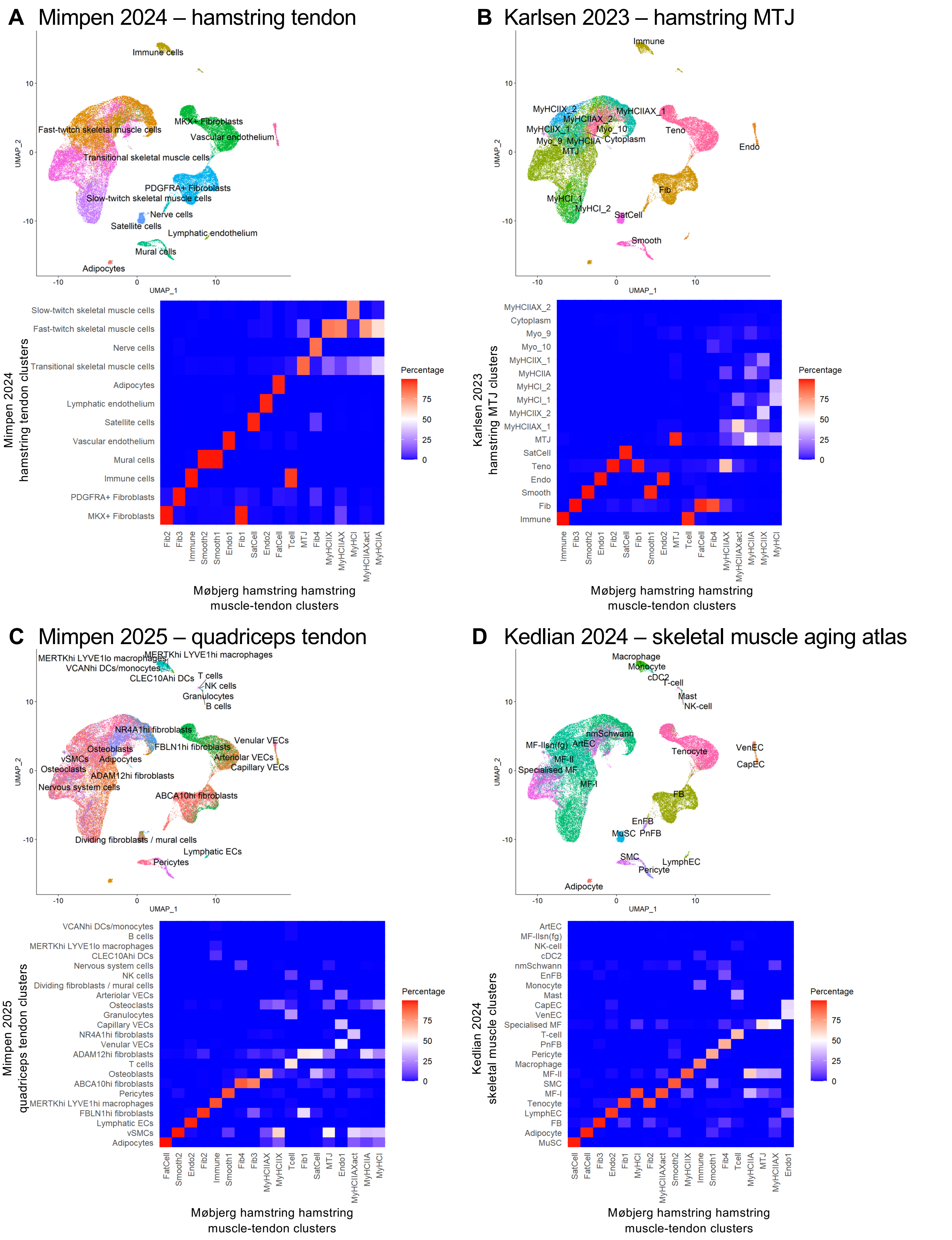


**Figure S5. Cross-dataset integration and correlation of fibroblast clusters.** Integration of the present study's data set with previously published snRNAseq and scRNAseq data sets: **(A)** healthy human hamstring tendon[25], **(B)** healthy human hamstring MTJ[23], **(C)** healthy and ruptured human quadriceps tendons[35], and **(D)** aging skeletal muscle atlas[36]. Cluster labels from published datasets were mapped onto the UMAP of the current study to enable cross-dataset comparison. Matrices display the percentage of nuclei in each cluster that align with corresponding clusters identified in published datasets.

### Møbjerg *et al.* – Supplementary Figure 6

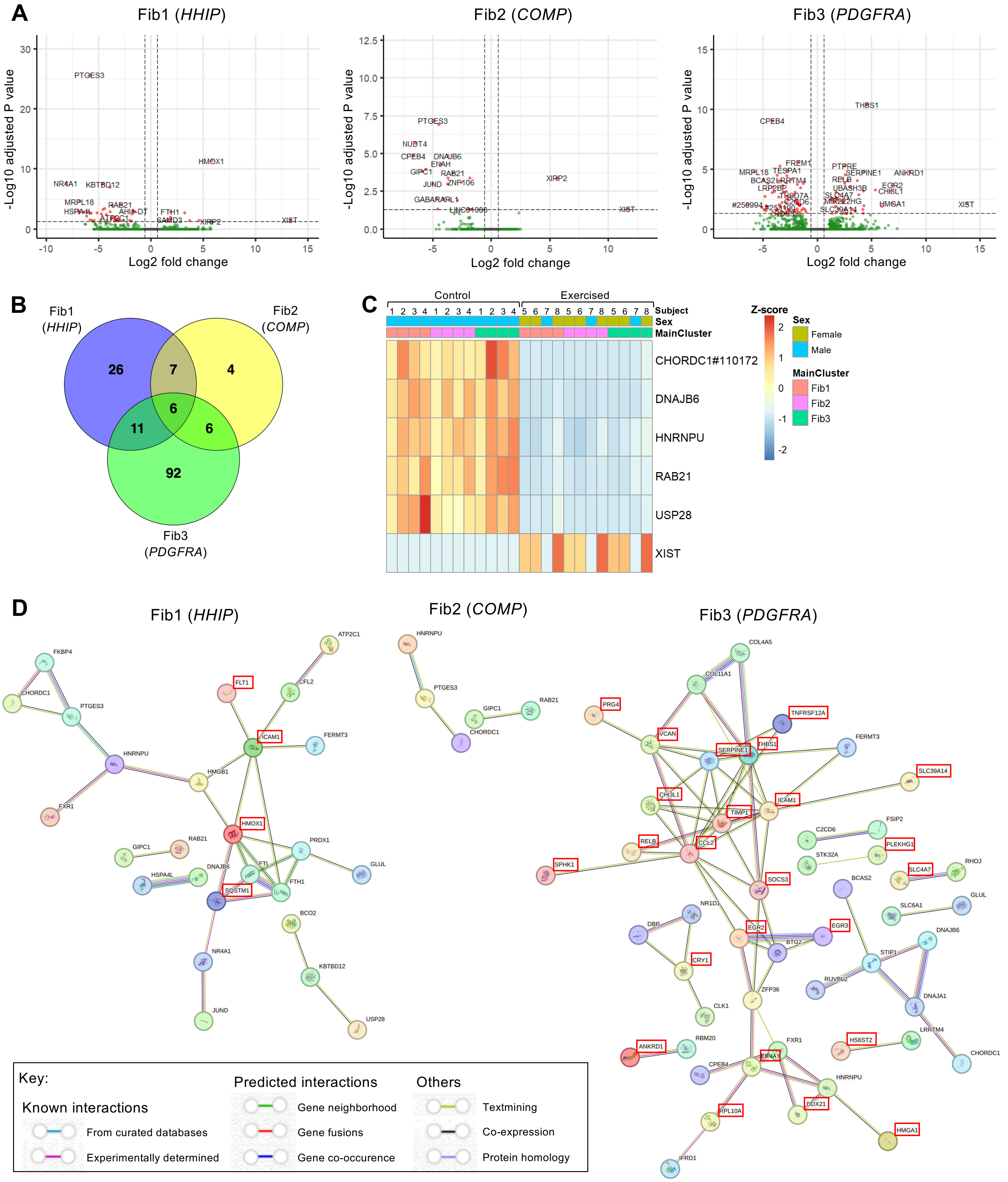


**Figure S6. Exercise-induced differentially regulated RNAs in fibroblast clusters. (A)** Volcano plots showing exercise-induced DEGs in Fib1, Fib2 and Fib3. No exercise regulated transcripts were detected in Fib4. Red data points indicate significant DEGs (log2 fold change>±0.585, FDR<0.05, DESeq2). Green data points indicate non-significantly changed RNAs. **(B)** Venn diagram showing number of common exercise-induced DEGs in the three fibroblast clusters. **(C)** Heatmap showing the expression levels of five commonly exercise-induced DEGs and XIST, the long non-coding RNA used in X-chromosome inactivation in females. Each column represents the average expression for a given replicate-cluster combination, annotated with biological sex. Gene expression values are row-scaled (z-score normalized) to show relative differences across samples for each gene. **(D)** Protein-protein interaction networks generated by STRING analysis for exercise regulated transcripts in Fib1, Fib2 and Fib3. Names outlined in red boxes indicate the gene was upregulated.

### Møbjerg *et al.* – Supplementary Figure 7

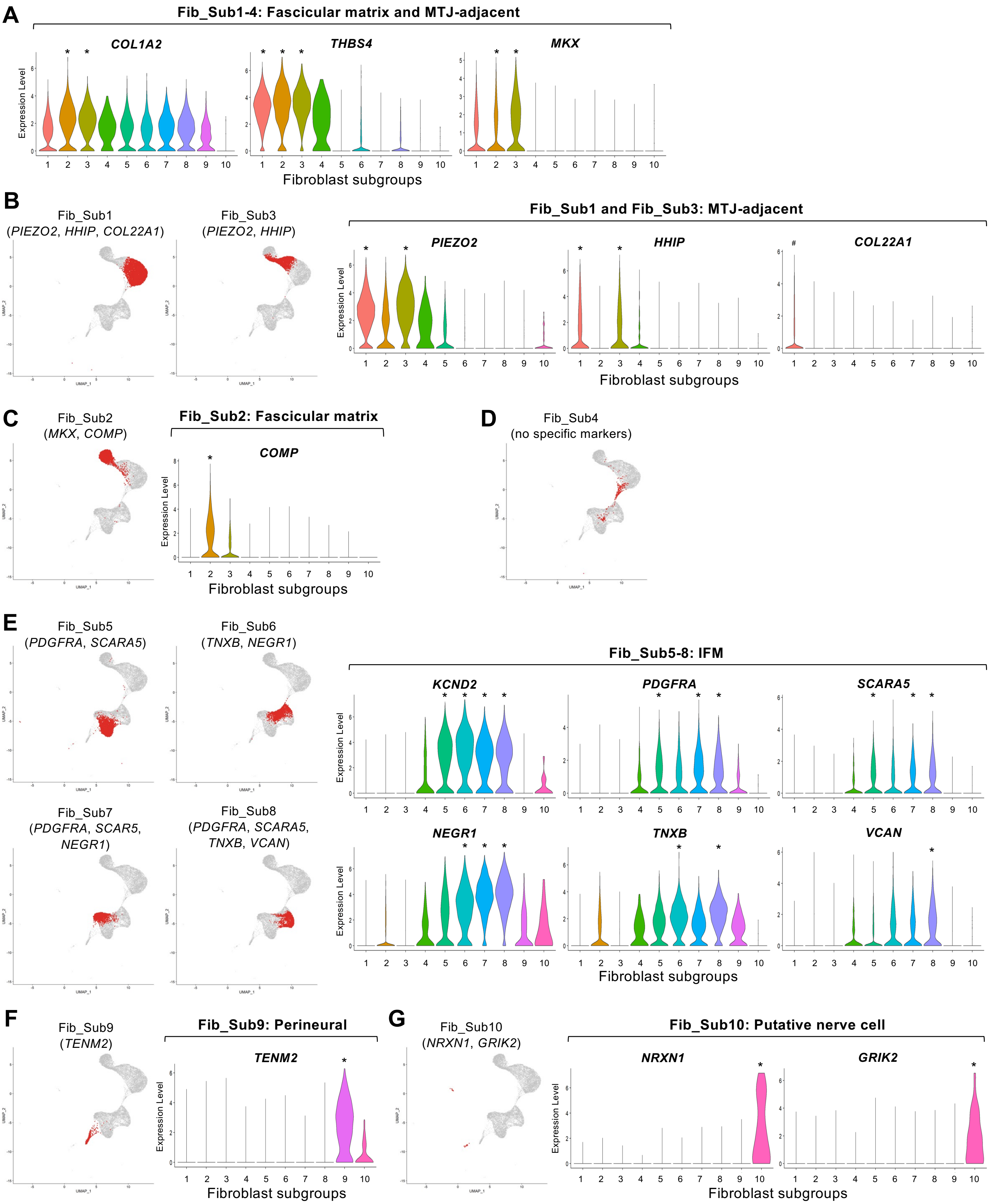


**Figure S7. Transcriptomic characterization of hamstring tendon fibroblast subclusters.** UMAPs show the distribution of each fibroblast subcluster in transcriptomic space, with accompanying violin plots highlighting key marker genes distinguishing each subcluster. (A) *COL1A2*, *THBS4*, and *MKX*. (B) Fib_Sub1 and Fib_Sub3 MTJ-adjacent: *PIEZO2*, *HHIP*, and *COL22A1*. (C) Fib_Sub2 fascicular matrix: *COMP*. (D) Fib_Sub4: no specific marker. (E) Fib_Sub5, Fib_Sub6, Fib_Sub7 and Fib_Sub8 IFM: *KCND2*, *PDGFRA*, *SCARA5*, *NEGR1*, *TNXB*, and *VCAN*. (F) Fib_Sub9 perineural: *TENM2*. (G) Fib_Sub10 putative nerve cell: *NRXN1* and *GRIK2*. * indicates significance (log2 fold change>1, FDR<0.05, DESeq) when compared to all clusters. # indicates significance (log2 fold change>1, FDR<0.05, DESeq) when compared within the fibroblast subclusters.

**Møbjerg *et al.* – Supplementary Figure 8**

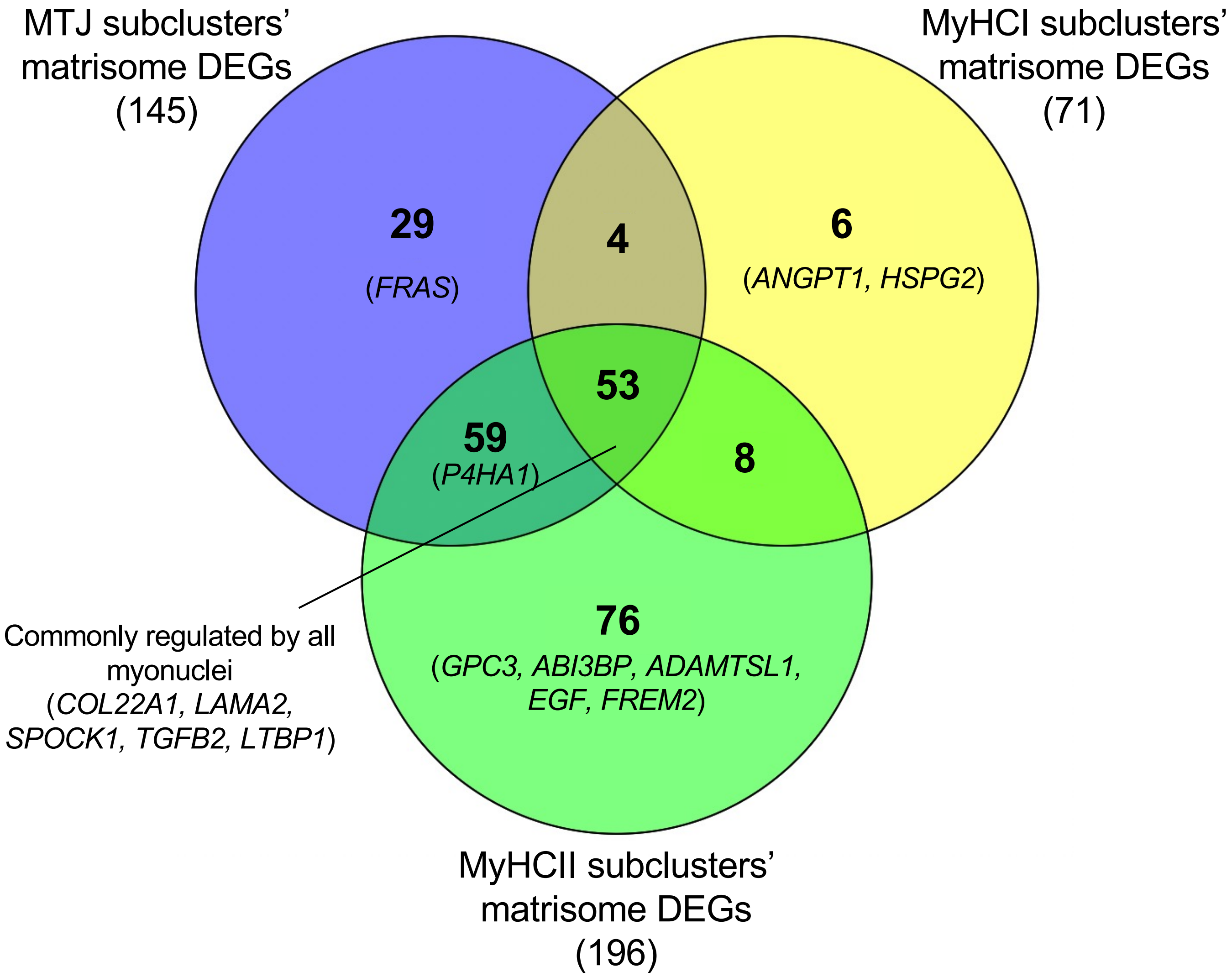


**Figure S8. Exercise-induced matrisome changes in myonuclear subtypes and their overlap with MTJ-enriched genes.** Venn diagram of exercise-induced DEGs (FDR<0.05, DESeq2; >±0.585 log2 fold change) identified in MTJ, MyHCI, and MyHCII subclusters. Genes are grouped according to their myonuclei identity, with duplicates across subclusters excluded. Gene names in parentheses are matrisome-annotated genes that are enriched in human hamstring MTJ[25].

### Møbjerg *et al.* – Supplementary Figure 9

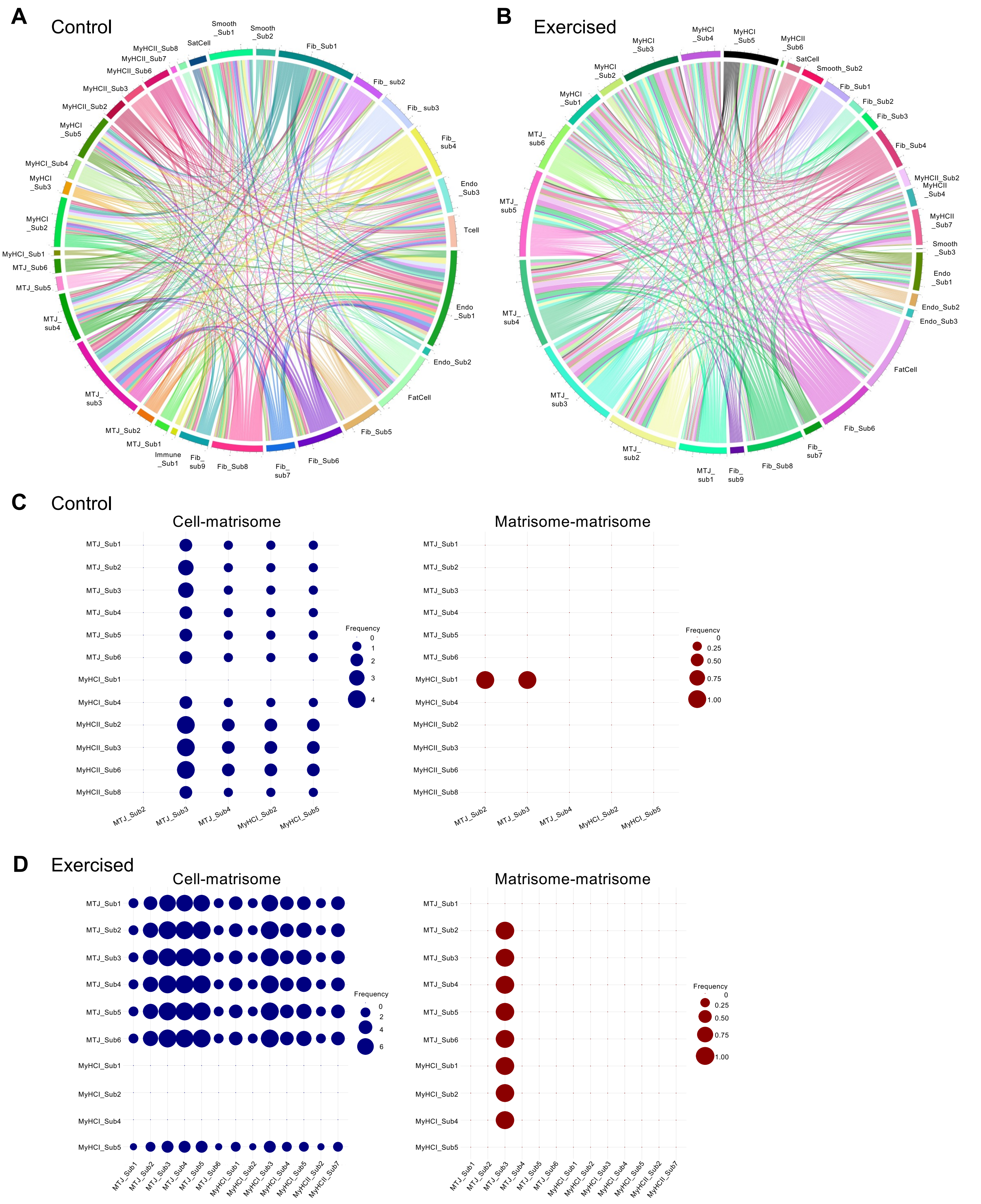


**Figure S9. Matrisome communication between nuclei subclusters in human hamstring muscle-tendon tissue. (A-B)** Chord diagrams show the distribution of cell-matrisome and matrisome-matrisome interactions between nuclei subclusters, including only interactions in which both partners are expressed in at least 80% of the nuclei participating in that interaction. Results are shown separately for **(A)** control and **(B)** exercised tissues. **(C-D)** Bubble plots showing frequency of cell-matrisome (left panels) and matrisome-matrisome (right panels) interactions between myonuclei subclusters in **(C)** control and **(D)** exercised tissues. Only interactions with the highest reliability score (level 3, MatriCom) were included in the bubble plots to ensure high stringency.
